# PKHD1L1 is required for stereocilia bundle maintenance, durable hearing function and resilience to noise exposure

**DOI:** 10.1101/2024.02.29.582786

**Authors:** Olga S. Strelkova, Richard T. Osgood, Chunjie J. Tian, Xinyuan Zhang, Evan Hale, Pedro De-la-Torre, Daniel M. Hathaway, Artur A. Indzhykulian

## Abstract

Sensory hair cells of the cochlea are essential for hearing, relying on the mechanosensitive stereocilia bundle at their apical pole for their function. Polycystic Kidney and Hepatic Disease 1-Like 1 (PKHD1L1) is a stereocilia protein required for normal hearing in mice, and for the formation of the transient stereocilia surface coat, expressed during early postnatal development. While the function of the stereocilia coat remains unclear, growing evidence supports PKHD1L1 as a human deafness gene. In this study we carry out in depth characterization of PKHD1L1 expression in mice during development and adulthood, analyze hair-cell bundle morphology and hearing function in aging PKHD1L1-defficient mouse lines, and assess their susceptibility to noise damage. Our findings reveal that PKHD1L1-deficient mice display no disruption to bundle cohesion or tectorial membrane attachment-crown formation during development. However, starting from 6 weeks of age, PKHD1L1-defficient mice display missing stereocilia and disruptions to bundle coherence. Both conditional and constitutive PKHD1L1 knock-out mice develop high-frequency hearing loss progressing to lower frequencies with age. Furthermore, PKHD1L1-deficient mice are susceptible to permanent hearing loss following moderate acoustic overexposure, which induces only temporary hearing threshold shifts in wild-type mice. These results suggest a role for PKHD1L1 in establishing robust sensory hair bundles during development, necessary for maintaining bundle cohesion and function in response to acoustic trauma and aging.

## Introduction

Auditory hair cells are highly specialized sensory cells located in the cochlea. Essential for our sense of hearing, they transduce the mechanical stimulus of sound into an electrical signal. Hair cells of the mammalian cochlea are terminally differentiated and do not regenerate once lost; therefore, hair-cell death or irreparable cochlear dysfunction leads to permanent sensorineural hearing loss. Hair-cell function is susceptible to disruption by genetic mutations and environmental factors such as noise exposure, ototoxic drug treatment, and aging.

We previously reported that the expression of Polycystic Kidney and Hepatic Disease 1-Like 1 (PKHD1L1), also known as fibrocystin like (Fibrocystin-L), is highly enriched in cochlear hair cells (Wu et al., 2019). PKHD1L1 is a very large, predominantly extracellular, membrane protein consisting of 4249 amino acids, with a single-pass transmembrane domain and a very short 6-amino acid intracellular cytoplasmic domain. PKHD1L1 has been identified as a biomarker or implicated in many cancers (Kang et al., 2023, 2023; Saravia et al., 2019; Song et al., 2021; L. Wang et al., 2023; Yang et al., 2023; Zheng et al., 2019; Zou et al., 2022); and has been linked to schizophrenia (Shang et al., 2024), and anxiety-like traits (Chitre et al., 2023).

There is increasing evidence supporting *PKHD1L1* as a human deafness gene. We previously showed that PKHD1L1-deficient mice develop delayed-onset progressive hearing loss (Wu et al., 2019). Similarly, knockout of *pkhd1l1* zebrafish paralogs, *pkhd1l1α* and *pkhd1l1β,* reduces auditory-evoked startle responses in zebrafish at larval stages, indicative of an early-onset hearing loss. (Makrogkikas et al., 2023). In humans, we recently reported biallelic variants of *PKHD1L1* in four unrelated families with congenital non-syndromic sensorineural hearing loss (Redfield et al., 2024). These PKHD1L1 variants were predicted to be damaging *in silico,* and some of the putative disease-causing mutations were shown to reduce the thermodynamic stabilities of PKHD1L1 protein fragments *in vitro* (Redfield et al., 2024). Furthermore, a large-scale exome study of 532 individuals recently identified the *PKHD1L1* gene as a possible contributor to hearing outcomes, with a higher variant load in older adults with hearing loss compared to the non-hearing loss group (Lewis et al., 2023). However, little is known about PKHD1L1 in the inner ear. In this study we aim to comprehensively characterize in mice the expression, localization, and roll of PKHD1L1 in hearing function and maintenance.

Within the cochlea there are two types of hair cells: one row of inner hair cells (IHC), responsible for sound transduction; and three rows of electromotile outer hair cells (OHC), which amplify sound-evoked vibrations within the cochlea. Hair cells derive their name from the actin-rich stereocilia they carry at their apical poles, which collectively form the mechanosensitive sensory hair bundle. Hair bundles are composed of three rows of stereocilia arranged in a staircase pattern. The mechanoelectrical-transducer (MET) machinery is located at the tips of the two shorter rows of stereocilia, linked to the adjacent taller row by fine protein filaments called tip links. Tip links mechanically gate the MET channels in response to bundle deflection, initiating the perception of sound and forming the basis of hearing (Beurg et al., 2009; Pickles et al., 1984). The tallest row of OHC stereocilia are embedded in the tectorial membrane (TM), a gelatinous extracellular matrix essential for hearing sensitivity and frequency selectivity (Legan et al., 2000, 2005; Sellon et al., 2019). The tips of the tallest row stereocilia are linked to the tectorial membrane by attachment crowns (Verpy et al., 2011). There are additional links between adjacent stereocilia, maintaining cohesion in the mature hair bundle. These inter-stereociliary links include horizontal top connectors in OHCs, and shaft connectors in IHCs (Goodyear et al., 2005; Nayak et al., 2007).

During the establishment of the mature hair bundle in early postnatal development there is a rapidly changing array of additional, temporary, links on the surface of the stereocilia. Identified by electron microscopy, these include transient lateral links, ankle links, and OHC shaft connectors; in addition to a dense surface coat on the stereocilia and apical membrane of the cell, visualized by staining with tannic acid (Goodyear et al., 2005). We previously found that PKHD1L1 localizes to the surface of stereocilia during early postnatal development. Using *Pkhd1l1^fl/fl^::Atoh1-Cre^+^* mice, we established that it is required for the formation of the stereocilia surface coat, especially towards the tip of stereocilia (Wu et al., 2019). We showed that *Pkhd1l1^fl/fl^::Atoh1-Cre^+^* mice lacking PKHD1L1 display no major morphological disruption to hair bundles at the onset of hearing and initially exhibit near-normal ABR and DPOAE thresholds, however, develop progressive hearing loss later in life (Wu et al., 2019). Here therefore, we investigate a case in which a developmental genetic perturbation, lacking an overt morphological or functional phenotype, can still cause serious hearing deficit in adulthood.

The functions of the stereocilia coat, and specifically the role of PKHD1L1 on the surface of stereocilia during development, is unclear. As the surface coat is only detected during the early postnatal development of cochlear hair cells, it has been suggested to play a role in bundle maturation (Goodyear et al., 2005). We previously hypothesized that PKHD1L1 could be required for the proper localization of other stereocilia components (Wu et al., 2019). The localization of PKHD1L1 and hearing loss phenotype previously observed in PKHD1L1-deficient mice suggest that PKHD1L1 may play a role in establishing connections between the OHC stereocilia and the tectorial membrane.

The sensory cells and structures of the cochlea, established during development, are not regenerated once lost. Hair cells must therefore survive a lifetime in order to maintain hearing function. In the present study, we provide evidence suggesting that PKHD1L1 is required for the establishment of robust sensory bundles, resilient to varied environmental stresses during aging. Experimentally, high noise levels can induce permanent (irreversible) threshold shifts (PTS), or temporary (reversible) threshold shifts (TTS) depending on exposure level and duration (Kujawa & Liberman, 2006; Ryan et al., 2016; Y. Wang et al., 2002). Administration of moderate noise insult in an experimental setting therefore allows us to assess the sensitivity of PKHD1L1-deficient hair cells to environmental insult compared to wildtype littermates.

In this study we expand our understanding of PKHD1L1 in the inner ear, and address the previously observed temporal disparity between PKHD1L1 expression and the onset of hearing loss in its absence. We comprehensively characterize *Pkhd1l1* gene and protein expression during development and in adult mice. Morphological analysis in PKHD1L1-deficient mice revealed no developmental disruption to bundle cohesion or attachment-crown formation. However, progressive bundle disruption and missing stereocilia are observed in mice deficient in PKHD1L1 from 6 weeks of age onwards, with concurrent high frequency hearing loss progressing to lower frequencies with age in both conditional and constitutive *Pkhd1l1* knock-out mice. We find that PKHD1L1-deficient mice have a higher susceptibility to permanent hearing loss following moderate acoustic overexposure. Here, therefore, we demonstrate a role for PKHD1L1 in establishing robust sensory hair bundles during development, which are required in later life to maintain function in response to acoustic insult and aging.

## Results

### *Pkhd1l1 mRNA* is expressed in outer and inner hair cells during development

Previously, we reported active transcription of *Pkhd1l1* in hair cells during early postnatal development (Wu et al., 2019). To investigate the discrepancy in the timeframe between the detection of PKHD1L1 protein and the physiological effects of its deletion on hearing function in later life, we first elected to directly, and more precisely, evaluate the *Pkhd1l1* mRNA levels within inner and outer hair cells during development and aging.

Using a commercially available probe targeting the 1061-2157 bp region (~8% of the 13049 bp transcript) of *Pkhd1l1* mRNA, we carried out *in-situ* hybridization by RNAscope in acutely dissected organ of Corti from P1 to 9-month-old mice. *Pkhd1l1* mRNA was visualized by fluorescence imaging in apical, middle, and basal turns of the cochlea. *Pkhd1l1* expression was observed as diffuse, as well as bright punctate labeling, in both IHCs and OHCs (Fig. 1a, and Sup. Fig. 1).

**Figure 1.**
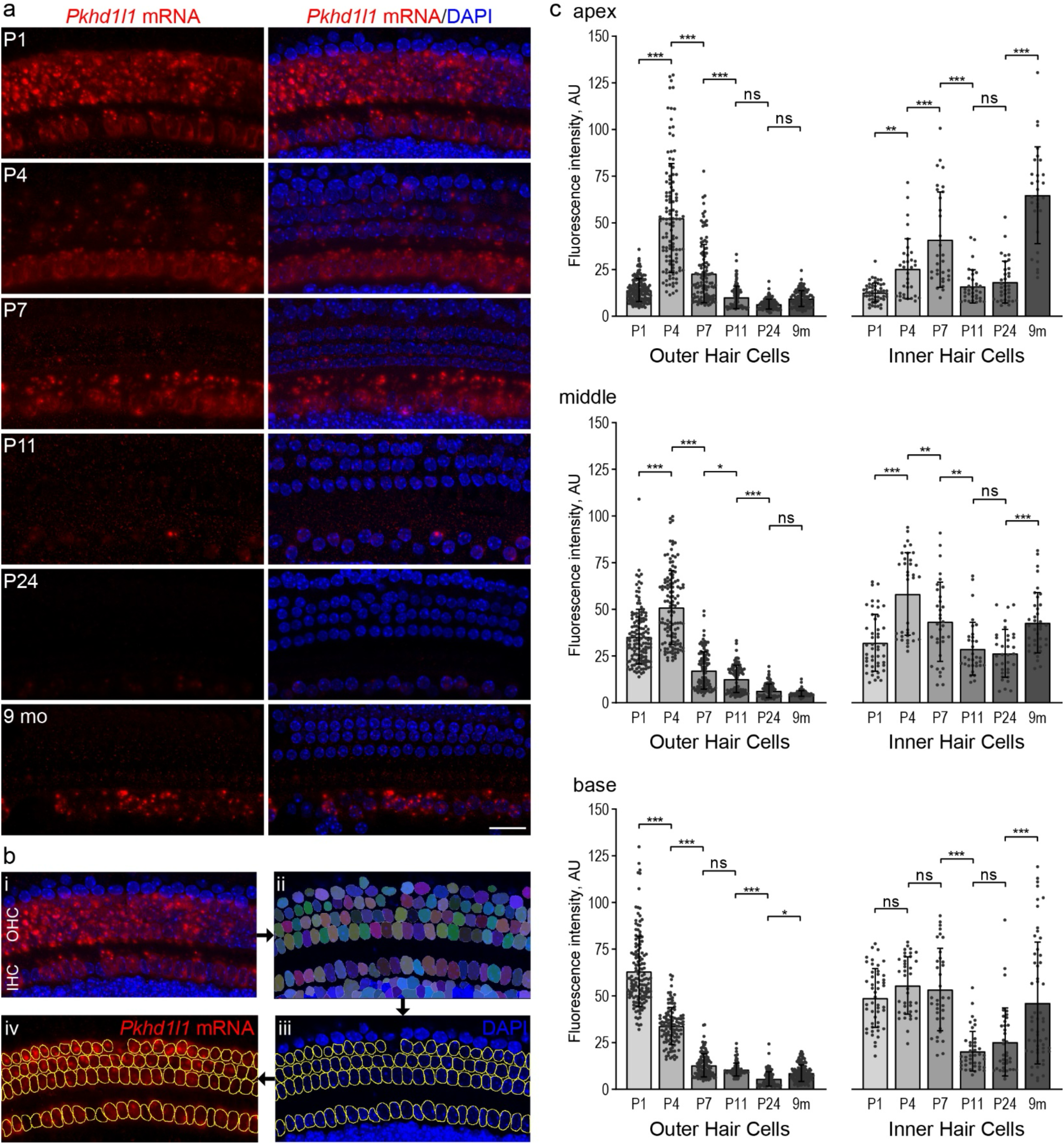
*Pkhd1l1* mRNA levels in hair cells gradually decrease during early postnatal development. **a**, *In situ* detection of *Pkhd1l1* mRNA by RNAscope fluorescence labeling of the basal cochlear region. Maximum intensity z-projections. *Red, Pkhd1l1* mRNA fluorescence. *Blue*, DAPI. *Scale bar,* 20 µm. **b**, Schematic of RNAscope single cell fluorescence intensity quantification approach: nuclei in all cells were individually segmented using *Cellpose* analysis pipeline based on DAPI label, the masks representing hair-cell nuclei were assigned to IHCs or OHCs to measure *Pkhd1l1* mRNA fluorescence labeling intensity on a single-cell level. **c**, Average *Pkhd1l1* mRNA fluorescence intensity measurements in three cochlear locations from P1 to 9 months of age in OHCs *(left)* and IHCs *(right)*. *Pkhd1l1* mRNA levels in OHCs significantly decrease by P24 in all cochlear regions and remain virtually undetectable. *Pkhd1l1* mRNA levels in IHCs show a similar gradual downregulation during development, however, increase in the 9-months-old cochlea. Points represent individual cells; bars show mean ± SD. Two cochleae were analyzed per condition (15-80 cells from each). *Statistical analysis*, one-way ANOVA with Sidak’s multiple comparison tests between adjacent time points, * p<0.05, ** p<0.01, *** p<0.001.

To quantify *Pkhd1l1* mRNA labeling intensity levels in individual cells, we utilized *Cellpose*, a deep-learning-based generalist algorithm for cellular segmentation (Stringer et al., 2021). Briefly, the DAPI-stained hair-cell nuclei were detected using *Cellpose,* and their ROIs transferred to imageJ for single-cell intensity measurements of mRNA labeling in maximum intensity z-projections (Fig. 1b).

The intensity of mRNA labeling varied in an age- and cochlear region-dependent manner. Considering first, OHCs: *Pkhd1l1* mRNA fluorescence labeling intensity is highest in the basal region at P1 and drops rapidly during early postnatal development (Fig. 1a & c). Peak mRNA expression in the middle and apical regions occurs later, at P4 (Fig. 1c & Sup. Fig. 1a). As hair cells towards the base of the cochlea develop before those at the apex, and peak fluorescence intensity is similar between the regions, differences in mRNA expression between regions at a given time point are likely developmental rather than tonotopic. In all 3 regions, OHC *Pkhd1l1* mRNA fluorescence intensity reduces to almost undetectable levels by P11 (comparable to negative control *dapB,* Sup. Fig. 1b), before the onset of hearing, and remains undetectable in mature hair cells at P24 and 9-month time points (Fig. 1 & Sup Fig. 1a).

Surprisingly, given that little PKHD1L1 was previously detected at the surface of IHCs at early postnatal time points (Wu et al., 2019), *Pkhd1l1* mRNA is also detected at comparatively high levels in inner hair cells. As with expression in OHCs, fluorescence intensity peaks first in early postnatal development (P7), although later than that observed in OHCs (P1-4). More curiously, however, *Pkhd1l1* mRNA is also detected at high levels in IHCs at the 9-month time point, (Fig. 1 & Sup. Fig. 1a, with no signal detected in *dapB* negative control, Sup. Fig. 1b). These findings suggest a possibility of PKHD1L1 protein expression in IHCs in the adult cochlea.

### PKHD1L1 is transiently expressed in cochlear hair cells during early postnatal development

The abovementioned *Pkhd1l1* mRNA expression labeling results do not correlate well in timing or location with our previous findings of protein detection (Wu et al., 2019). Therefore, to more fully characterize the expression pattern of PKHD1L1, we carried out anti-PKHD1L1 immunofluorescence labeling in the cochlea throughout development and in adulthood (Fig. 2). Labeling was carried out in P4–9 month old *Pkhd1l1^fl/fl^::Atoh1-Cre^−^* mice (Fig. 2 top 7 panels), facilitating the use of previously reported *Pkhd1l1^fl/fl^::Atoh1-Cre^+^* mice as a negative labeling control (Fig. 2, bottom panel).

**Figure 2.**
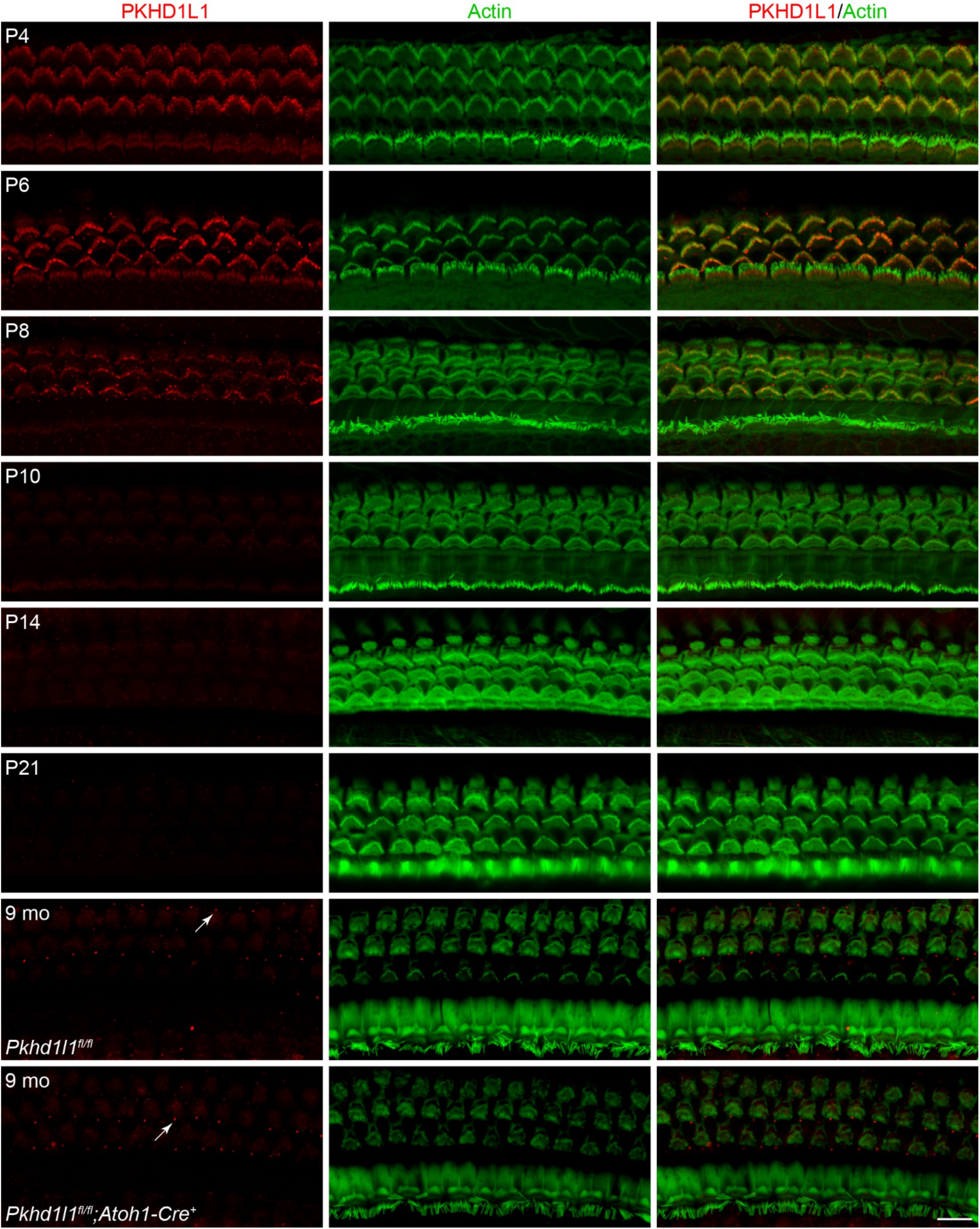
PKHD1L1 is only detected in hair cells during early postnatal development. Anti-PKHD1L1 immunolabeling reports presence of PKHD1L1 protein in stereocilia bundles of early postnatal mice, with labeling intensity gradually decreasing by P10-P14. No specific anti-PKHD1L1 fluorescence labeling was observed in adult cochleae at 9 months, while some non-specific labeling is evident within basal bodies of kinocilia and primary cilia of support cells (white arrows). The *bottom* panel is from *Pkhd1l1^fl/fl^::Atoh1-Cre^+^* mouse (negative control), while all other panels are from normal, *Pkhd1l1^fl/fl^::Atoh1-Cre^−^* mice. *Red*, anti-PKHD1L1; *Green*, phalloidin labeling. Micrographs are maximum intensity z-projections of hair-cell apical poles and stereocilia bundles. Images are representative of >6 cochleae per time point. *Scale bar,* 10 µm.

In agreement with our previous report, PKHD1L1 was detected in hair cells throughout the first postnatal week (Fig. 2 P4-P8) along the length of the cochlea (Sup. Fig. 2a). PKHD1L1 labeling was observed on the hair bundle stereocilia, with no specific signal detected in hair cell bodies (Sup Fig. 2c). Anti-PKHD1L1 labeling intensity gradually decreased with development, was largely gone by P10, and completely cleared in mature hair cells at P21.

Although PKHD1L1 is detected on inner hair cell bundles at P4 and P6, despite largely similar *Pkhd1l1* mRNA expression results between IHCs and OHCs, protein labeling intensity was higher at the surface of the OHC stereocilia bundles compared to IHCs (Fig. 2 compared to Fig. 1A). Furthermore, no anti-PKHD1L1 immunoreactivity was observed in IHC at 9 months. Fluorescence signal detected within the basal bodies of kinocilia and primary cilia of support cells in *Pkhd1l1^fl/fl^::Atoh1-Cre^−^* mice at this time point, is also found in *Pkhd1l1^fl/fl^::Atoh1-Cre^+^* cochlea, devoid of PKHD1L1, (Fig. 2 arrows, comparing bottom 2 panels) and is therefore concluded to be non-specific. Similarly, no specific anti-PKHD1L1 immunolabeling signal was detected in the mouse utricle at P0 and P4 (data not shown).

From these data, we conclude there is not a direct correlation between in *Pkhd1l1* mRNA levels and protein expression levels detected using immunolabeling techniques. PKHD1L1 protein expression labeling is restricted to early postnatal development and localizes to the hair bundles of both IHCs and OHCs.

### PKHD1L1 expression is not redistributed on OHC hair bundle stereocilia during development

During early postnatal development, a transient surface coat, and several types of transient stereocilia links are present on the developing stereocilia bundle (Goodyear et al., 2005). PKHD1L1 was previously demonstrated to localize with the surface coat at P4 (Wu et al., 2019). Several key stereocilia components including the tip-link proteins cadherin-23 (CDH23) (Michel et al., 2005) and protocadherin-15 (PCDH15) (Indzhykulian et al., 2013; Kazmierczak et al., 2007); horizontal top connector and attachment crown protein stereocilin (STRC)(Verpy et al., 2011); and transient ankle link component Vlgr1 (Goodyear & Richardson, 1999), have been demonstrated to be spatially refined, concentrated, or redistributed, following their initially more diffuse expression on the developing bundle. To determine the distribution pattern of PKHDlL1 on OHC bundles during this developmental period, scanning electron microscopy (SEM) immunogold labeling of PKHD1L1 was carried out on cochleae between P4 - P10 (Fig. 3).

**Figure 3.**
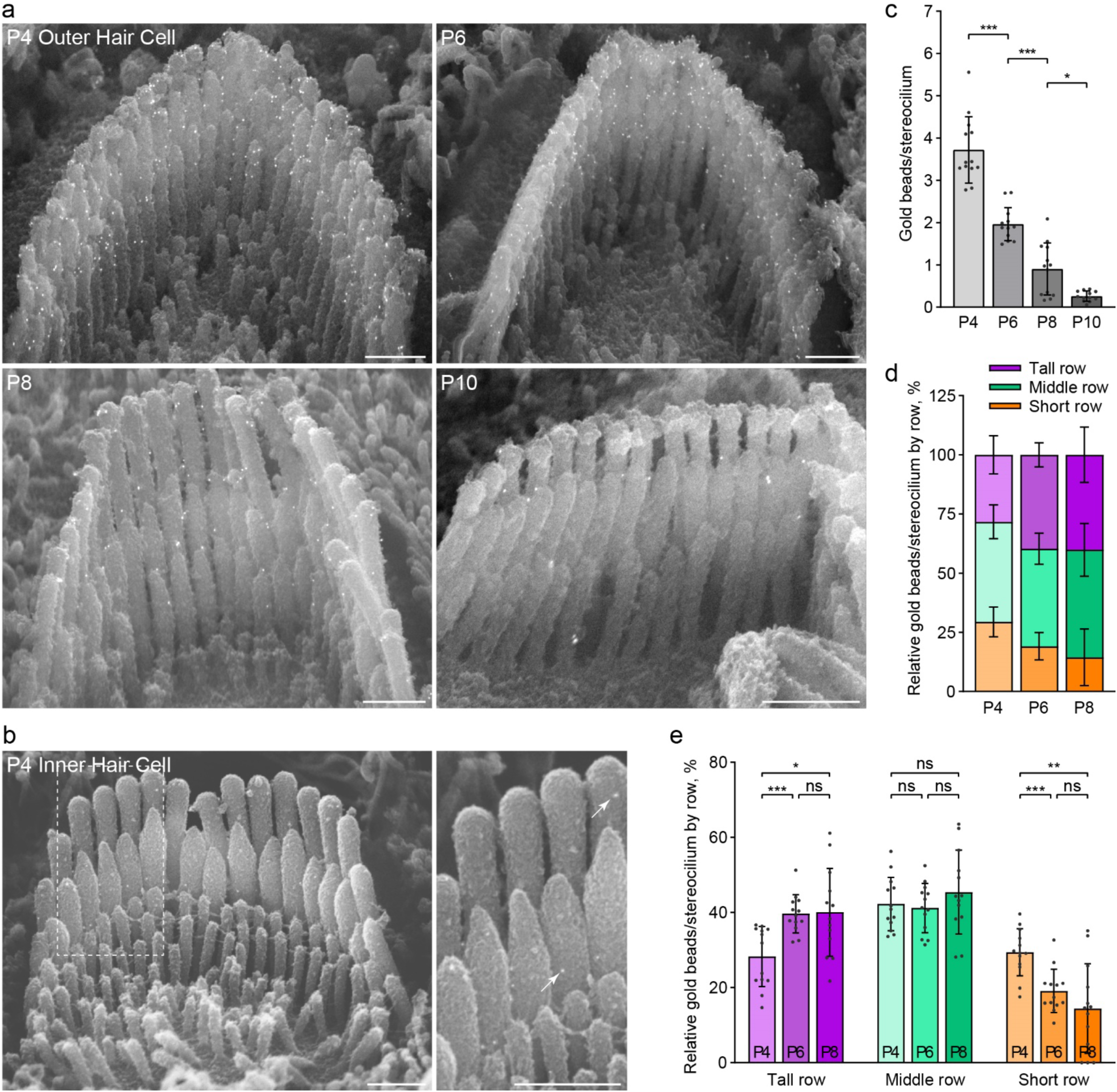
The distribution of anti-PKHD1L1 immunogold labels on the surface of OHC bundles does not change during development. **a**, Anti-PKHD1L1 immunogold SEM of OHC bundles in the mid/basal region at P4, P6, P8 and P10. **b**, P4 IHC bundles are labeled with fewer gold beads (arrows), as compared to OHCs. *Scale bars,* 500 nm. **c**, Quantification of the total number of gold beads per OHC stereocilium show a gradual decrease of labeling density, with most of the labeling gone by P10. *Statistical analysis*, one-way ANOVA with Sidak’s multiple comparison tests between consecutive time points. **d**, **e,** Distribution of gold beads at the surface of the tallest, middle, and short rows of stereocilia as a portion of the total number of gold beads per bundle, normalized per stereocilium. A slight increase in the proportion of gold beads in the tallest row, and a reduction in the proportion of gold beads in the shortest row, is observed between P4 and P6. *Statistical analysis*, two-way ANOVA with matched design between stereocilia rows from the same cell. Tukey’s multiple comparison tests are shown between time points for each row. * p<0.05, ** p<0.01, *** p<0.001. All results are presented as mean ± SD, datapoints represent individual cells (n=13 at each time point).

Since EDTA decalcification is required for tissue dissection to enable immunogold SEM analysis in cochleae older than P6, we first evaluated the effect of EDTA treatment on immunogold labeling efficiency. There was no significant difference in the number of gold particles observed on stereocilia bundles in P4 mouse cochleae, processed with and without decalcification buffer (Sup. Fig. 3).

In agreement with immunofluorescence data (Fig 2), immunogold labeling on OHC bundles was observed between P4–P10. Labeling is greatest at P4 with almost no labeling observed by P10 (Fig. 3). Very little gold labeling was detected in IHCs, unlike that observed in immunofluorescence (Fig. 3b compared to Fig. 2.). This is likely as a result of reduced labeling efficiency of the immunogold technique.

Upon close inspection of gold beads at the surface of stereocilia bundles, PKHD1L1 did not appear to localize to any specific areas or links prior to the cessation of its expression at P10 (Fig. 3a). In agreement with previously reported data at P4 (Wu et al., 2019), PKHD1L1 is found on all three stereocilia rows throughout development. To evaluate the distribution of PKHD1L1 on specific stereocilia rows of OHC bundles during its expression, we quantified the proportion of gold particles located on the tallest, middle and short stereocilia rows at each time point (Fig. 3d,e). There is a slight increase in the proportion of gold beads located on the tallest row, and decrease on the shortest row, between P4 and P6. However, this does not represent a substantive change in the overall distribution of PKHD1L1 between the stereocilia rows. We therefore conclude that PKHD1L1 is not spatially refined or redistributed on the surface of OHC hair bundles during development.

### PKHD1L1 contains potential proteolytic cleavage sites

To reconcile our findings of *Pkhd1l1* mRNA expression in adult IHC with the lack of antibody labeling of the PKHD1L1 protein, we looked more closely at the structure of PKHD1L1 and antibody epitope (Fig. 4). We previously reported that PKHD1L1 labelling, unlike tip links and horizontal top connectors, is sensitive to treatment with protease subtilisin (R. J. Goodyear et al., 2005; Wu et al., 2019). To investigate the possibility of PKHD1L1 cleavage further we carried out cleavage site prediction on the protein sequence of *Pkhd1l1* orthologs from 10 species (Sup. Table 1). Using the ProP 1.0 online prediction tool (Duckert et al., 2004), we identified three potential cleavage sites in the PKHD1L1 sequence (numbered based upon the mouse sequence): p.R3566, p.R3976, and p.R3986 (Sup. Fig. 4).

**Figure 4.**
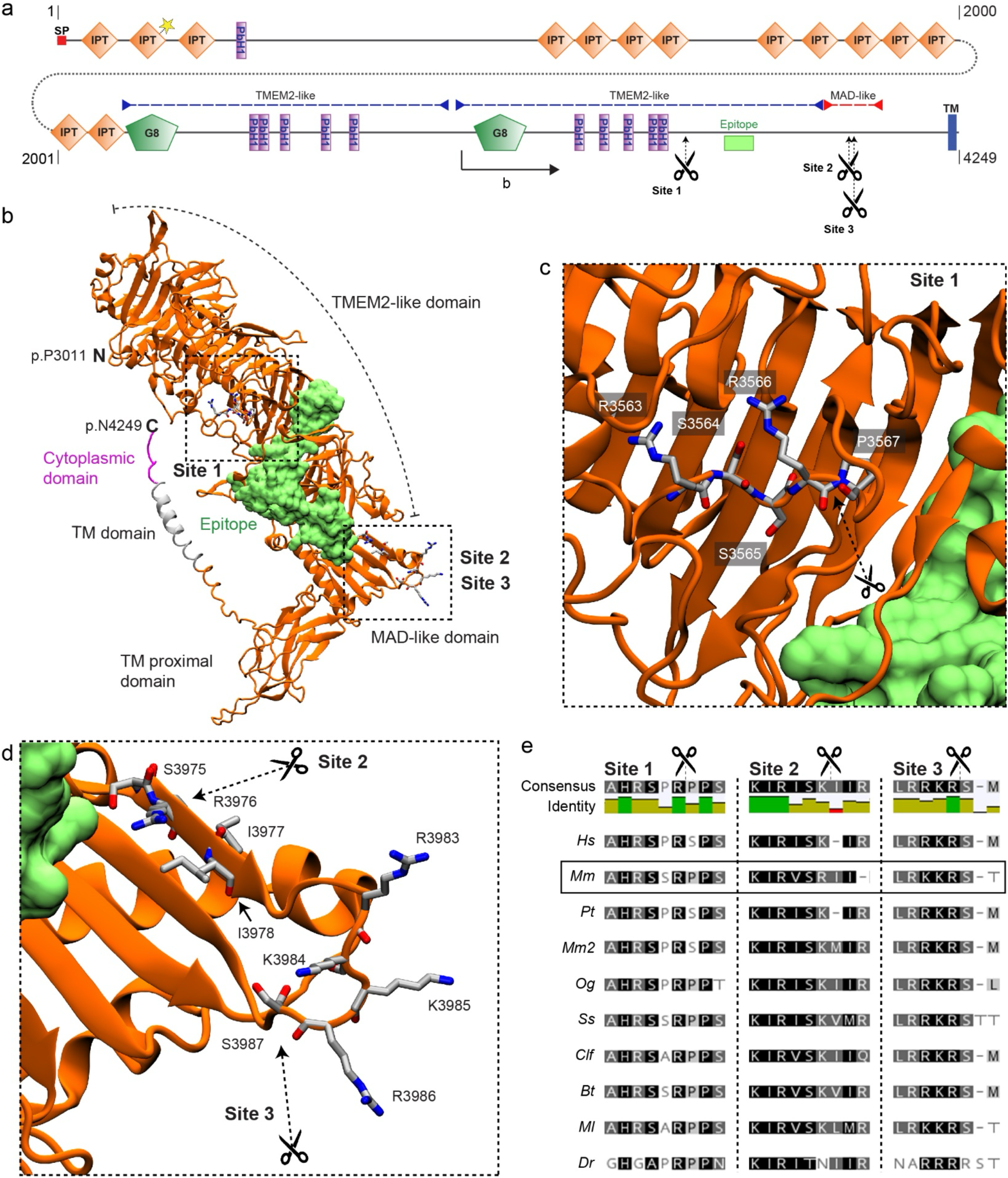
PKHD1L1 contains potential proteolytic cleavage sites. **a**, PKHD1L1 domain diagram, based on consensus of UniProt and SMART predictions (domain predictions in Sup. Table 2). Residue numbering includes the signal peptide (20 amino acids) using the mouse reference sequence NP_619615.2. Position of frame shift and premature stop codon in PKHD1L1-deficient mice is depicted by dotted line and yellow star. Purple dashed lines indicate TMEM2-like domains, red dashed line indicates MAD-like domain. The epitope for anti-PKHD1L1 antibody used in this study (NBP2-13765) is depicted by a lime box. Potential cleavage sites at p.R3566 (site 1), p.R3976 (site 2), and p.R3986 (site 3) are depicted with scissors. **b**, AlphaFold2 modelling was used to predict the structure of a *Mm* PKHD1L1 protein fragment comprising the residues p.P3011-4249 (illustrated on panel a by black arrow). The protein fragment is depicted in cartoon representation of the ectodomain fragment (orange), transmembrane domain (grey) and cytoplasmic domain (pink). The epitope for anti-PKHD1L1 antibody NBP2-13765 —against p.3670-p.3739, is depicted in surface representation (lime). Potential cleavage are depicted with scissors. The antibody epitope region and the transmembrane domain are separated by the PKHD1L1 MAD-like domain with predicted cleavage sites 2 and 3, suggesting that labeling of the PKHD1L1 ectodomain might be absent following PKHD1L1 cleavage. **c, d,** Insets representing zoomed-in views of the putative cleavage sites. Images were rendered with the VMD software. **e**, Protein sequence alignment of the predicted PKHD1L1 cleavage site regions across 10 different species (see accession codes in Sup. Table 1). Analysis was carried out with the Geneious software. Background colors represent the rate of residue conservation: black indicates high-sequence similarity, while white indicates poor residue similarity or its absence. A consensus sequence is shown for all the species at the top of the panel, with a color-coded identity map to illustrate the level of consensus between the different species.

We utilized Alphafold2 to model the structure of a PKHD1L1 fragment (p.3011-p.4249) and mapped the potential cleavage sites (Fig. 4a-d). Site 1 (p.R3566), with a confidence score below threshold in mice but present in other species, is located in a parallel beta-helix repeat (PbH1) rich region with structural homology to the human TMEM2 protein ectodomain (Fig. 4a-c). While site 2 (p.R3976), not identified in other species, and with a confidence score below threshold, and site 3 (p.R3986, with the highest confidence score in all species) are located within a domain with structural homology to the PCDH15 MAD12 domain (Fig. 4a,b & d). Notably, cleavage site 3 is highly conserved across 10 different orthologs and lies within an exposed loop of PKHD1L1 MAD-like domain (Fig. 4e). Proteolytic cleavage at site 3 would release an extracellular protein fragment containing the NBP2-13765 antibody epitope (Fig. 4 lime), thus hindering the labeling of the remaining membrane-tethered portion of the PKHD1L1 peptide by the NBP2-13765 antibody used for immunolabeling in this study. The remaining membrane-bound peptide would have an extracellular domain of just 236 amino acids.

Although detected *Pkhd1l1* mRNA and protein levels correlate well during early-postnatal development, this analysis suggests that protein cleavage could explain the discrepancy in mRNA detection and protein labeling in the IHCs of 9-month-old mice.

### Tectorial membrane attachment crowns and imprints retain normal morphology in PKHD1L1-deficient mice

The progressive hearing loss phenotype we previously observed in PKHD1L1-deficient mice (Wu et al., 2019) is similar to that observed in mice with tectorial membrane deficits, such as in STRC and CEACAM16-deficient mice (Goodyear et al., 2019; Verpy et al., 2008). Interaction with the tectorial membrane, established during early postnatal development, is critical for OHC function. We therefore sought to investigate the establishment, and maintenance of OHC attachment with the tectorial membrane in PKHD1L1-deficient mice as a possible source of pathology.

STRC plays a key role in attachment crown formation between OHC tall row stereocilia and tectorial membrane (Verpy et al., 2011). We detected STRC at the tips of stereocilia in *Pkhd1l1^fl/fl^::Atoh1-Cre^+^* mice, as in *Pkhd1l1^fl/fl^* littermate controls, by immunofluorescence at P35 – a time point when bundles and the tectorial membrane are fully developed (Fig. 5a,b). Immunogold SEM of bundles demonstrates the correct localization of STRC, primarily in halos around the tip of the tallest stereocilia, in addition to between stereocilia tips, in both PKHD1L1-deficient mice and littermate controls (Fig. 5c). We therefore conclude that PKHD1L1 is not required for the localization of STRC to attachment crowns or horizontal top connectors. Furthermore, OHC bundle imprints in the tectorial membrane, observed by SEM, still form in PKHD1L1-defficient mice, and were still present after 6 months (Fig. 5d). Together these data suggest that the establishment of OHC links with the TM are not disrupted in PKHD1L1-defficient mice.

**Figure 5.**
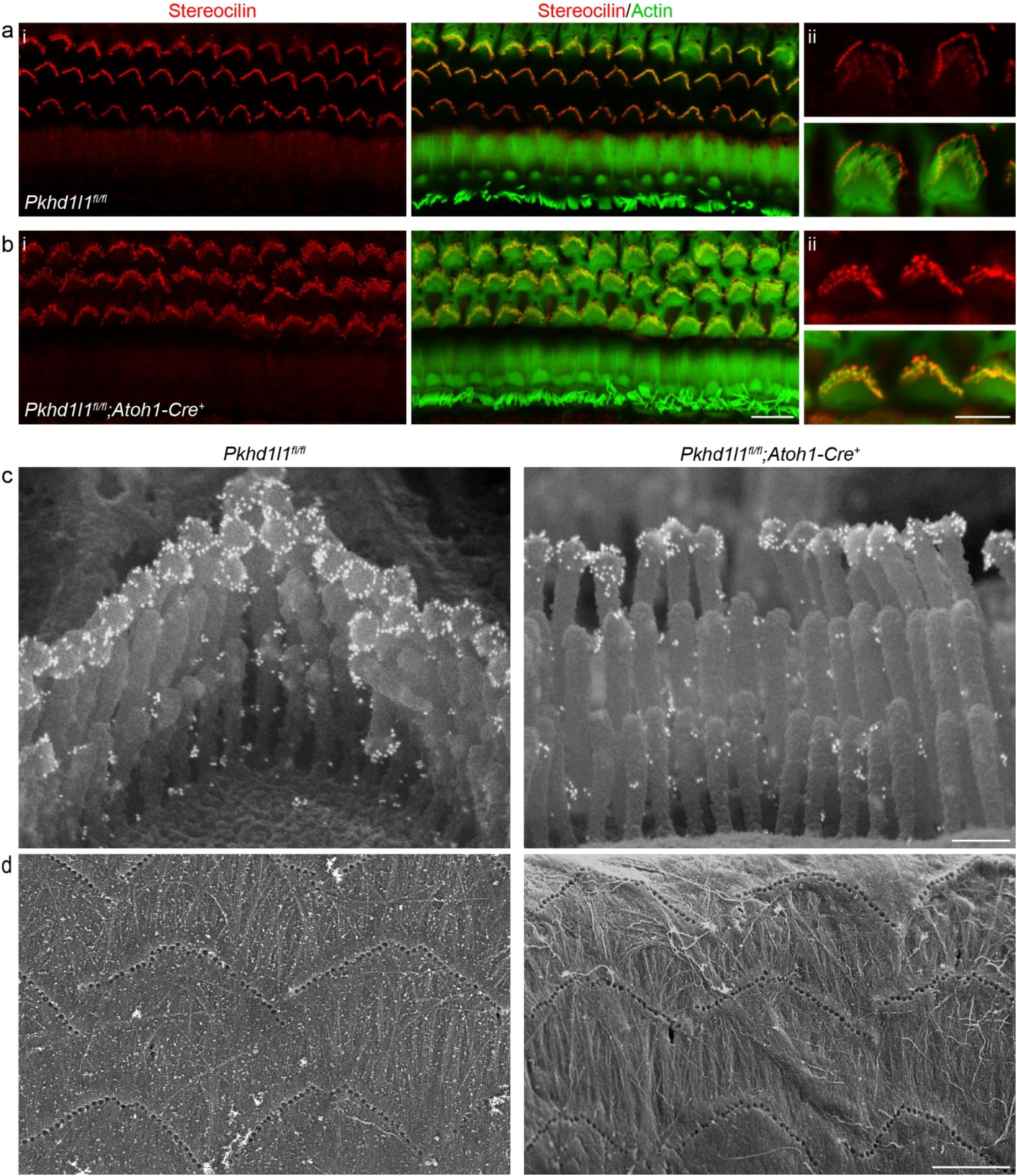
Stereocilin is properly localized to the OHC stereocilia bundles in PKHD1L1-deficient mice. Anti-STRC immunofluorescence labeling of OHCs in the basal region of P35 mouse cochlea localizes to the tips of stereocilia (**a**) *Pkhd1l1^fl/fl^::Atoh1-Cre^−^* controls and (**b**) *Pkhd1l1^fl/fl^::Atoh1-Cre^+^* mice (ii, inset with higher magnification). *Red*, anti-PKHD1L1; *Green*, phalloidin labeling. Images are representative of 2 cochleae per genotype. **c,** Anti-STRC immunogold SEM of OHC stereocilia in *Pkhd1l1^fl/fl^::Atoh1-Cre^+^* and their *Pkhd1l1^fl/fl^::Atoh1-Cre^−^* control mice at P29. The gold bead labels form a well-characterized ring pattern at the tips of tall row stereocilia in control and PKHD1L1-deficient mice. Images are representative of 2 cochleae per genotype. **d,** SEM imaging of TM reveal normal imprints in 6-month-old *Pkhd1l1^fl/fl^::Atoh1-Cre^+^* mice and *Pkhd1l1^fl/fl^::Atoh1-Cre^−^* controls. Images are representative of 3 cochleae per genotype. *Scale bars*, *a-b* i, 10 µm; ii, 5 µm; *c*, 250 nm; *d,* 5 µm.

### Loss of PKHD1L1 induces late-onset progressive hearing loss and stereocilia disorganization

Despite the restricted temporal expression of PKHD1L1 to early postnatal development, our previous study revealed progressive hearing loss in PKHD1L1^−^ deficient mice, tested up to 6 months. However, no gross morphological changes were found during adolescence (Wu et al., 2019). We sought to further evaluate the progression of hearing loss in *Pkhd1l1^fl/fl^::Atoh1-Cre^+^* conditional hair-cell-specific knock out (cKO) and constitutive *Pkhd1l^−/−^* KO mice as they age, in addition to conducting comprehensive stereocilia analysis by SEM in PKHD1L1-deficient mice during aging.

The *Atoh1-Cre^+^* line expresses Cre as early as embryonic day 10.5. In juveniles Cre expression, and in turn *Pkhd1l1* KO in *Pkhd1l1^fl/fl^::Atoh1-Cre^+^* mice, is limited to sensory hair cells, as well as some supporting cells and cochlear and vestibular sensory neurons (Matei et al., 2005). Hearing deficit was observed by ABR recordings in cKO *Pkhd1l1^fl/fl^::Atoh1-Cre^+^* mice compared to *Pkhd1l1^fl/fl^* (Cre^−^) littermates (Fig. 6a). ABR thresholds were elevated at 6 weeks in the high-frequency region (32 kHz, 45 kHz), progressing to lower-frequency regions (16 kHz, 22 kHz) by 9 months of age. Consistent with the predominant localization of PKHD1L1 to OHC bundles, hearing deficits were mirrored in DPOAE thresholds. Thresholds were elevated first at 32 kHz at 6 weeks, progressing in severity and frequency range by 9 months (Fig. 6a), indicative of progressive OHC dysfunction.

**Figure 6.**
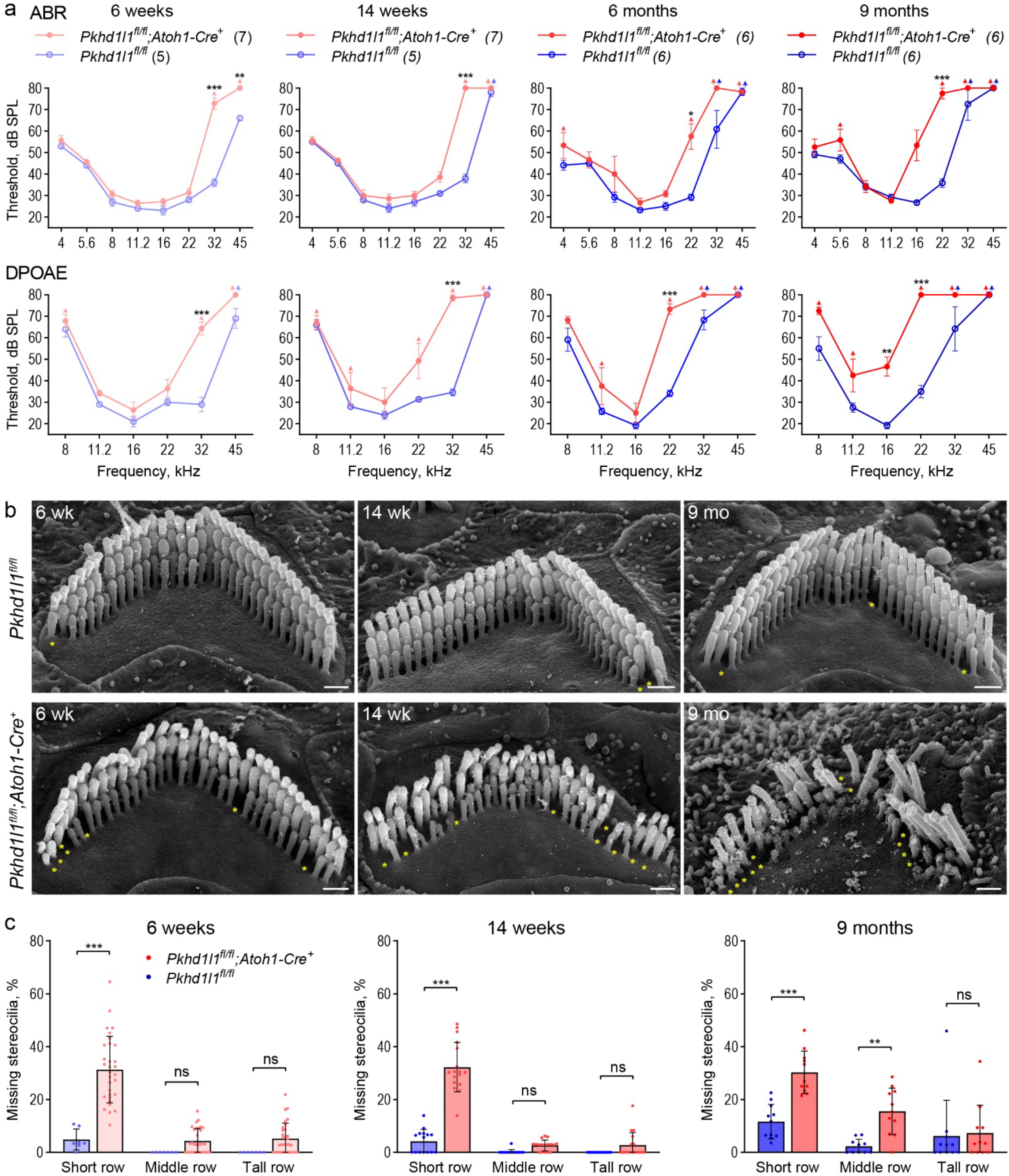
PKHD1L1-deficient mice display progressive hearing loss and stereocilia disruption. **a**, ABR and DPOAE of *Pkhd1l1^fl/fl^::Atoh1-Cre^+^* and *Pkhd1l1^fl/fl^::Atoh1-Cre^−^* mice show progressive high-frequency hearing loss in PKHD1L1-deficient mice as early as 6 weeks. Data are shown as mean ± SEM. Up-arrows indicate that at the highest SPL level tested (80 dB SPL), at least one mouse in the group had no detectable thresholds at that frequency. *Statistical analysis*, two-way ANOVA with matched design between frequencies from the same mouse. Sidak’s multiple comparison tests are shown between genotypes for each frequency. **b**, SEM of OHC stereocilia bundles at 6 weeks, 14 weeks, and 9 months in the basal region (32 kHz) reveal progressive stereocilia loss (asterisks) and bundle disorganization. *Scale bars*, 500 nm. **c**, Quantification of missing stereocilia in OHC bundles in the basal turn (32 kHz region) of the cochlea, presented as percentage of stereocilia missing per row. There is a greater stereocilia loss in *Pkhd1l1^fl/fl^::Atoh1-Cre^+^* mice as compared to their *Pkhd1l1^fl/fl^::Atoh1-Cre^−^* littermates at all age points. Data displayed as mean ± SD, points represent individual cells (see n in *Methods*). *Statistical analysis*, two-way ANOVA with matched design between rows from the same cell. Sidak’s multiple comparison tests are shown between rows for each genotype. *p<0.05, **p<0.01, ***p<0.001.

The use of hair-cell-specific *Pkhd1l1* KO in *Pkhd1l1^fl/fl^::Atoh1-Cre^+^* mice is advantageous to mitigate against phenotypes related to possible unknown developmental or central processing functions of PKHD1L1 in other tissues and cell types. However, elevated ABR thresholds have been reported in mice with hair cells expressing Cre (Matern et al., 2017). In order to confirm that hearing deficit was not as a result of Cre expression, we generated constitutive KO *Pkhd1l^−/−^* mice. Constitutive KO of PKHD1L1 was not embryonically lethal; *Pkhd1l^−/−^* mice appear to develop normally and breed well. Elevated ABR and DPOAE thresholds were found in *Pkhd1l^−/−^* mice from 12 weeks, comparable to those observed in *Pkhd1l1^fl/fl^::Atoh1-Cre^+^* mice, again progressing to lower frequencies with age, tested up to 15 months (Sup. Fig. 5). These data confirm progressive hearing deficits as a result of PKHD1L1 KO.

The morphology of OHC stereocilia bundles was examined using SEM in *Pkhd1l1^fl/fl^::Atoh1-Cre^+^* cKO hair cells during the progression of hearing deficit (Fig. 6b). Mild stereocilia loss (Fig. 6b, yellow asterisks) was observed after 6 weeks, becoming more pronounced over time. To analyze the progression of OHC stereocilia loss from as a result of aging, we calculated the number of missing stereocilia from each row of the bundle. Notably, stereocilia loss was most significant in the shortest row, progressing to the taller rows with age; most prominently in, but not limited to, the high frequency basal region (Fig. 6c - 32 kHz region, Sup. Fig. 6 - 22 kHz and 16 kHz regions).

Hair cell stereocilia bundles of *Pkhd1l1^fl/fl^::Atoh1-Cre^+^* cKO mice at 6 weeks of age show no overt signs of disrupted development or disorganization, as compared to *Pkhd1l1^fl/fl^* controls (Fig. 6b). However, bundle disorganization is observed at 14 weeks increasing in severity by 9 months of age. OHC bundle structure was well maintained in *Pkhd1l1^fl/fl^* littermate controls, with rare instances of stereocilia loss. These data implicate a role for PKHD1L1 expression during development in the maintenance and potential stabilization of stereocilia bundles, and in turn OHC function, in later life.

### Acoustic overexposure leads to permanent hearing loss in PKHD1L1-dificent mice

Moderate noise trauma paradigms have been shown to induce temporary threshold shifts (TTS) in hearing function, which are subsequently recovered to normal levels (Jensen et al., 2015; Y. Wang et al., 2002). Despite the expression of PKHD1L1 being restricted to early postnatal development, PKHD1L1-deficient mice display delayed onset progressive phenotypes in ABRs, DPOAEs and OHC bundle morphology (Fig. 6 & Sup. Fig. 5). This could be indicative of a protein required for the establishment of more durable hair bundle morphology and function. To test this hypothesis, we assayed bundle and hearing heath and durability following noise overexposure.

Exposure to 94 dB SPL 8-16 kHz bandpass noise for 2 hours was carried out at 6 weeks of age in *Pkhd1l ^fl/fl^::Atoh1-Cre^+^* mice and *Pkhd1l1^fl/fl^* littermate controls – a time point and frequency range at which there is no hearing deficit in *Pkhd1l1^fl/fl^::Atoh1-Cre^+^* mice (Fig. 6a). Higher noise levels were initially trialed in wild-type mice, however these lead to permanent threshold shifts (PTS), and were therefore considered too damaging for BL6 background mice to recover from.

ABRs and DPOAEs were measured at 1 day, 2 days, 2 weeks, and 8 weeks, after noise exposure and compared to baseline levels collected prior to the noise insult, in order to determine threshold shifts resulting from the noise exposure (Fig. 7a). In both *Pkhd1l1^fl/fl^* and *Pkhd1l1^fl/fl^::Atoh1-Cre^+^* mice ABR and DPOAE thresholds were elevated in the 16-32 kHz region 1 day post noise exposure with some recovery after 2 days (Fig. 7b). Thresholds recovered back to baseline levels by 2 weeks post-noise exposure in *Pkhd1l1^fl/fl^* mice, consistent with TTS. Thresholds did not recover to baseline levels in *Pkhd1l1^fl/fl^::Atoh1-Cre^+^* mice however, with threshold shifts maintained at 2 and 8 weeks following noise insult (Fig. 7b).

**Figure 7.**
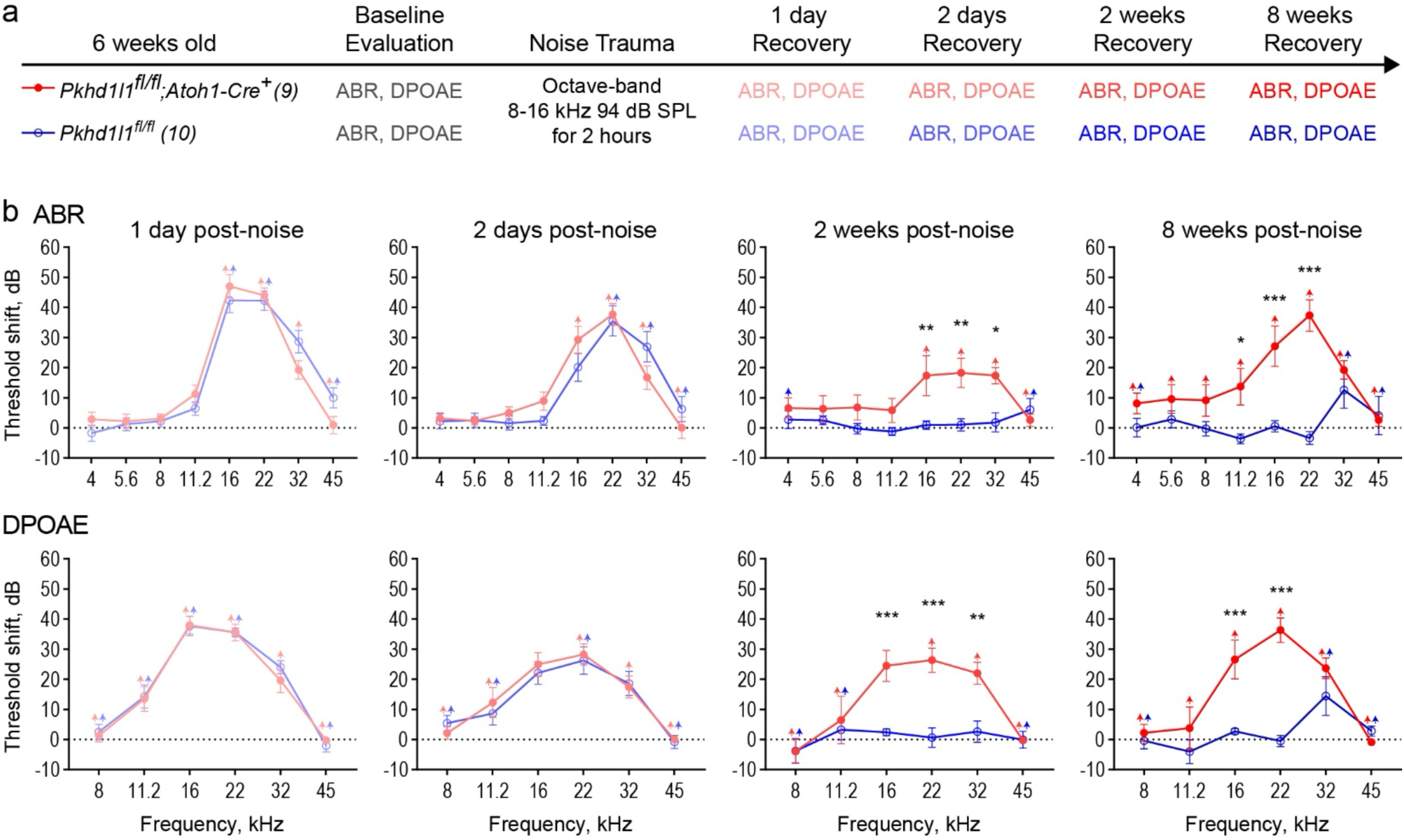
PKHD1L1-deficient mice experience permanent hearing loss after noise trauma levels that induce TTS in control animals. **a**, Experimental design. 6-week-old *Pkhd1l1^fl/fl^::Atoh1-Cre^+^* mice and *Pkhd1l1^fl/fl^::Atoh1-Cre^−^* control animals had baseline ABR and DPOAE evaluation prior to TTS-inducing level of noise trauma (8-16 kHz, 94 dB SPL). ABR and DPOAEs were subsequently recorded at 1 day, 2 days, 2 weeks, and 8 weeks, post noise exposure. **b**, ABR and DPOAE threshold shifts from pre-noise baseline evaluation. *Pkhd1l1^fl/fl^::Atoh1-Cre^−^* control mice (*blue*) exhibit temporary threshold shifts in ABRs and DPOAEs followed by recovery to baseline levels 2 weeks post-noise trauma, while *Pkhd1l1^fl/fl^::Atoh1-Cre^+^* mice (*red*) show only slight recovery 2 days after noise trauma, and permanent threshold shift in ABRs and DPOAEs at later time points. Data are shown as mean ± SEM. Up-arrows indicate that at the highest SPL level tested, at least one mouse in the group had no detectable thresholds at that frequency prior to calculation of threshold shift. *Statistical analysis*, two-way ANOVA with matched design between frequencies from the same mouse. Sidak’s multiple comparison tests are shown between genotypes for each frequency. *p<0.05, **p<0.01, ***p<0.001.

In the absence of noise trauma (Sup. Fig. 7) threshold shifts of <10 dB were observed at high frequencies between the 6-week evaluation time point and an additional 8-week ‘recovery’ time point in both genotypes, as a result of aging in these mice. This does not however account for the large, pan-frequency, threshold shifts observed in PKHD1L1-deficient mice alone following noise exposure. These data therefore show the noise exposure, normally only sufficient to cause TTS in normal mice, leads to PTS in PKHD1L1-deficient mice.

We examined OHC bundle morphology in *Pkhd1l1^fl/fl^::Atoh1-Cre^+^* mice and *Pkhd1l1^fl/fl^* littermates 2 and 8 weeks following noise exposure, time points at which TTS were recovered in control mice but not PKHD1L1-defficent mice (Fig. 8). In control *Pkhd1l1^fl/fl^* mice, bundles remained intact and well-organized at 2 and 8 weeks post-noise exposure. In contrast, in *Pkhd1l1^fl/fl^::Atoh1-Cre^+^* mice, bundle disorganization was observed 2 weeks following noise exposure (Fig. 8a), particularly in tallest row stereocilia, in addition to some loss of short row stereocilia, as expected at this age in PKHD1L1-defficient mice (comparison to Fig. 6b). By 8 weeks post-noise exposure, bundles were further disrupted, with instances of tall row stereocilia loss and disorganization (Fig. 8b). These findings demonstrate that deficiency in PKHD1L1 increases susceptibility to bundle disruption during moderate acoustic overexposure, leading to permanent threshold shifts.

**Figure 8.**
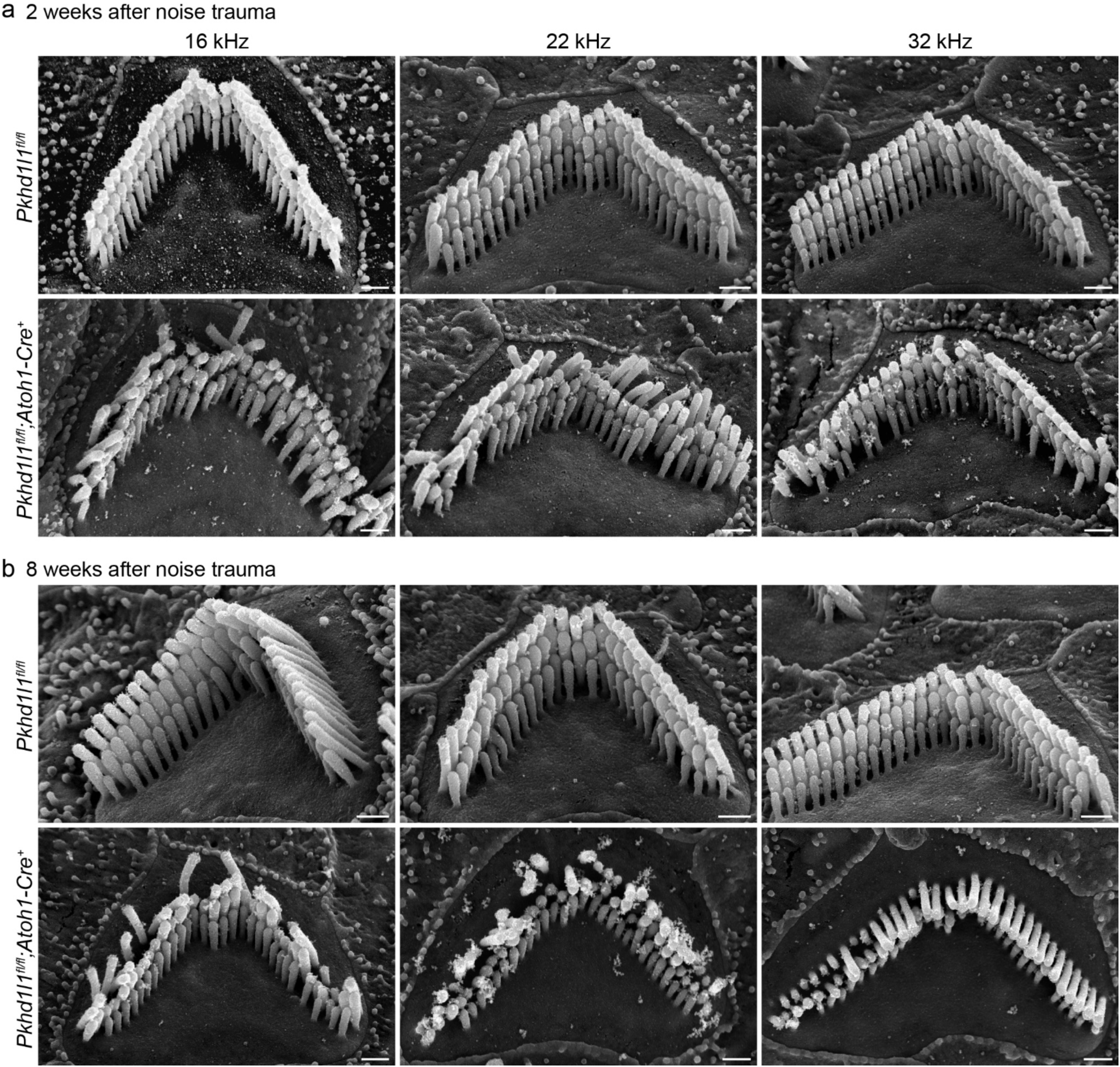
PKHD1L1-deficient mice exhibit stereocilia loss and bundle disorganization following moderate acoustic overexposure. SEM of OHCs following TTS-level noise trauma. **a,** 2 weeks after noise exposure *Pkhd1l1^fl/fl^::Atoh1-Cre^+^* bundle morphology is disrupted in comparison with *Pkhd1l1^fl/fl^::Atoh1-Cre^−^* littermate controls, also exposed to noise. **b,** 8 weeks after noise trauma stereocilia bundles in *Pkhd1l1^fl/fl^::Atoh1-Cre^+^* mice are further disrupted with instances of missing and shortened tall row stereocilia. *Scale bar*, 500 nm.

## Discussion

In this study we aim to better understand the role of PKHD1L1 in the sensory hair bundle and hearing function. Genetic perturbation of *Pkhd1l1* does not cause overt morphological and functional disruption during development, however, still causes serious hearing-related deficit in adulthood.

Here, we first fully characterized *Pkhd1l1* gene and protein expression during development and in adult mice. We confirmed developmentally restricted expression of PKHD1L1 on the sensory hair bundles of inner and outer hair cells. We further demonstrated by immunogold SEM that PKHD1L1 does not localize or redistribute to specific stereocilia bundle structures during this developmental period. Furthermore, we observe progressive bundle disruption and missing stereocilia in mice deficient in PKHD1L1 from 6 weeks of age onwards. We report hearing threshold shifts, initially restricted to high frequencies, subsequently progressing to lower frequencies, in both conditional hair-cell specific and constitutive *Pkhd1l1* KO-out mice. Finally, we show that PKHD1L1-deficient mice have an increased susceptibility to permanent hearing loss following moderate acoustic overexposure, implicating the expression of PKHD1L1 during development in the establishment of stable bundles that are robust to environmental insult in later life.

### PKHD1L1 expression, localization, and potential cleavage

PKHD1L1 protein was detected during the first 10 days of postnatal hair bundle development, primarily localized on the stereocilia of OHC, with some expression observed in IHC (Fig. 2&3, and Sup. Fig. 2). Gene expression analysis using RNAscope revealed corresponding early-postnatal *Pkhd1l1* gene expression. However, unexpectedly, transcription of *Pkhd1l1* was also observed in the IHCs of 9-month-old adult mice (Fig. 1) despite the absence of corresponding protein detection (Fig. 2). This discrepancy suggests that PKHD1L1 protein detection could be absent due to lack of mRNA translation in IHC at this time point, or that the translated protein was not detectable at the IHC stereocilia by immunolabeling.

PKHD1L1 is predicted, *in silico,* to have a very short cytoplasmic domain consisting of only 6 amino acids, a single-pass transmembrane domain, and a very large 4202 amino acid extracellular domain (Fig. 4 models C-terminal residues p.3011-p.4249). In our previous study we demonstrated that PKHD1L1 labeling (by antibody NBP2-13765, also used in this study, with an epitope between p.3670-p.3739 Fig. 4a lime) is sensitive to the protease subtilisin (Wu et al., 2019). Furthermore, PKHD1L1 has been identified in the ‘sheddome’ of activated blood platelets by mass spectrometry (Fong et al., 2011). The presence of PKHD1L1 following activation was substantially reduced by inhibitors of the protease ADAM17. These reports suggest that PKHD1L1 is susceptible to proteolytic cleavage *in vitro* and *in vivo*.

Here, we identified potential proteolytic cleavage sites in PKHD1L1 by *in silico* analysis across *Pkhd1l1* orthologs. R3986 is highly conserved across 10 PKHD1L1 orthologs, and scores highly with the ProP cleavage site prediction tool (Duckert et al., 2004) (Sup. Fig. 4). This cleavage site is located in the MAD-like domain, with structural homology to the PCDH15 MAD12 domain (De-la-Torre et al., 2018). Referred elsewhere as the SEA domain, this domain occurs in many cell surface proteins, in which it undergoes proteolytic cleavage or autoproteolysis(Anderson et al., 2024; Pei & Grishin, 2017). Cleavage at R3986 would release an extracellular protein fragment containing the NBP2-13765 antibody epitope, preventing detection of the remaining 263 amino acid PKHD1L1 fragment with this antibody. Therefore, PKHD1L1 protein cleavage could explain the discrepancy in *Pkhd1l1* mRNA detection and protein labeling in the IHCs of 9-month-old mice, if *Pkhd1l1* mRNA is translated at this time point. In addition, PKHD1L1 splice variants lacking the NBP2-13765 epitope region in adult IHCs cannot be ruled out. However, to our knowledge, there have been no verified alternate PKHD1L1 isoforms reported in humans or mice in publicly accessible databases.

*Pkhd1l1* mRNA (Fig. 1) and protein (Fig. 2) detection in early development correlate well and cease before P24 and P21 timepoints, respectively. We have not observed PKHD1L1 labeling on the tectorial membrane of young mice (P6-P10, data not shown), which might be expected upon PKHD1L1 cleavage. As hearing threshold shifts have been reported at as early as 3 weeks of age in PKHD1L1-dfficent mice (Wu et al., 2019), prior to the unexplained mRNA detection in IHCs, we believe the early postnatal expression of PKHD1L1 is likely to be responsible for its primary function observed in this study, and more specifically the phenotypic effects observed during its deficit. Nonetheless, whether PKHD1L1 is cleaved *in vivo* in adult mouse cochleae, and the potential role of the cleaved PKHD1L1 fragments, is an interesting area for further study.

### The role of PKHD1L1 and the stereocilia coat

The function of the stereocilia coat during bundle maturation in early postnatal development remains unclear. Although often considered together, distinct from the developmentally transient surface coat labeled with tannic acid, the stereocilia and apical surface of hair cells are covered by a glycocalyx coat, which can be visualized by cationic dyes such as Alcian blue (Santi & Anderson, 1987). This glycocalyx coat persists in mature hair cells, and has been suggested to contribute to bundle cohesion during stimulation and/or preventing stereocilia fusion (Dolgobrodov et al., 2000a, 2000b; Karavitaki & Corey, 2010). However, PKHD1L1 expression is limited to early postnatal development (Fig. 2 & 3) and is primarily localized to the external face of the stereocilia rather than between them (Wu et al., 2019), suggesting a distinct role.

Deficits in key mechanotransduction machinery or stereocilia link proteins often cause morphological disruption of the hair bundle during development, resulting in a profound hearing loss (reviewed by Petit and Richardson (Richardson & Petit, 2019)). In contrast, PKHD1L1-deficient mice undergo normal bundle maturation (Fig. 2 & 3), and exhibit only high-frequency hearing loss at younger ages (Fig. 6 & Sup. Fig. 5). This indicates that PKHD1L1 expression during development is likely not essential for the gross maturation of the hair bundle, or the localization and function of the MET complex.

We previously proposed that PKHD1L1 may play a role in establishing connection between the OHC stereocilia with the tectorial membrane. The late-onset mild hearing loss phenotype, and increased DPOAE thresholds, observed in PKHD1L1-deficient mice is similar to that of previously reported in CEACAM16-deficient mice with TM deficits (Goodyear et al., 2019). Furthermore, PKHD1L1 is primarily located on OHC, towards the tips of the stereocilia (Fig. 3)(Wu et al., 2019), where TM attachment crowns form. Here, however, we present data that suggests the role of PKHD1L1 is neither (1) critical to, or (2) limited to, TM attachment crown formation.

1. PKHD1L1 is not critical for attachment crown formation. We observed normal localization of key attachment crown protein stereocilin in PKHD1L1-defficient mice (Fig. 5). Minor disruptions to bundle coherence are observed during development in animals with mutated attachment crown proteins stereocilin, otogelin, otogelin-like or tubby (Avan et al., 2019; Han et al., 2020). Similar disruption is also observed in PKHD1L1-difficient mice, however after initially normal development. Furthermore, unlike in mice deficient in attachment crown proteins (Avan et al., 2019; Han et al., 2020), stereocilia imprints are still present in the tectorial membrane of PKHD1L1-defficient mice (Fig. 5d). TM imprints formed during development are likely permanent however (Morisaki et al., 1991), therefore these data do not preclude the possibility of TM attachments weakening in PKHD1L1-defficient mice as they age and/or are exposed to noise insult.
2. PKHD1L1 function is likely not limited to attachment crown formation. PKHD1L1 is observed on the surface of IHCs (Fig. 2), which do not form attachment with the TM. Attachment crowns only form on the tallest row of OHC stereocilia, and as such stereocilin redistributes to OHC tall row stereocilia during development (Verpy et al., 2011). PKHD1L1, on the other hand, is expressed on all stereocilia rows and no major redistribution or refinement in the localization of PKHD1L1 to the tallest row of stereocilia is observed during its expression period (Fig. 3). Finally, stereocilia loss is observed in PKHD1L1-deficient mice, initially primarily in short-row stereocilia (Fig. 6b & c). This phenotype is unlikely to be caused by disruption of tall row stereocilia attachment to the TM.

PKHD1L1-difficent mice are sensitive to moderate acoustic overexposure (Fig. 7). Curiously, following noise damage, stereocilia loss and disorganization in PHD1L1-deficient mice was observed in tall row stereocilia (Fig. 8). Tall row damage after noise exposure in PKHD1L1-deficient mice could result from aberrant interaction between the stereocilia and tectorial membrane, or an inability of the bundle to cope with the potential increased strain on the stereocilia during noise trauma. Intriguingly in the *Tubby* mutant *tub/tub;Map1a^AKR^* mice, attachment crowns are present in the absence of horizontal top connectors. These mice are deficient in the attachment crown and horizontal top connector component TUB, combined with the restorative *Map1a* polymorphism on an AKR/N background. They display normal DPOAEs, while also exhibiting an increased sensitivity to noise damage (Han et al., 2020). As PKHD1L1-difficient mice display elevated DPOAE thresholds (Fig. 6 and Sup. Fig. 5) in addition to increased sensitivity to noise damage (Fig. 7), the loss of horizontal top connectors alone would not account for the phenotypes observed in these mice. However, the role of PKHD1L1 in the maintenance of stereocilia links, beyond attachment crowns, established during early postnatal development and often important for bundle cohesion (Goodyear et al., 2005), is worth further investigation.

### PKHD1L1 is required for the formation of robust stereocilia bundles in noise trauma and aging

As hair cells in the cochlea cannot regenerate once lost, the development of cells and structures resilient to damage during a lifetime, and mechanisms to absorb and repair damage, are essential for hearing longevity. Moderate acoustic overexposure induces temporary shifts in hearing thresholds (Wang et al., 2002). Upon recovery, normal mice exhibit no disruption to hair bundle morphology (observed Fig. 7 *Pkhd1l1^fl/fl^* controls), however, TTS-levels of acoustic overexposure can induce other lasting neuronal pathologies outside the scope of this study (Fernandez et al., 2020; Kujawa & Liberman, 2009; Liberman & Dodds, 1987). In the absence of PKHD1L1, threshold shifts in response to moderate noise insult are not reversed, concurrent with morphological disruption of stereocilia bundles (Fig. 7 & 8). We propose that PKHD1L1 expression during early postnatal bundle maturation is required for the establishment of robust sensory bundles, resilient to damage and disruption in response to noise exposure and aging.

We previously reported elevated ABR and DPOAE thresholds in *Pkhd1l1^fl/fl^::Atoh1-Cre^+^* mice as early as 3 weeks of age in high frequency regions, progressing across all tested frequencies by 6 months (Wu et al., 2019). In the present study we again report progressive hearing loss, initially at high frequencies, however, elevations in ABR and DPOAE thresholds were less pronounced and progressed to lower frequencies more slowly than observed previously (Fig. 6). We further verified these findings by generating and evaluating the hearing function in constitutive *Pkhd1l^−/−^* KO mice, in which a similar phenotype is observed (Sup. Fig. 5).

The ABR and DPOAE recordings in these two studies were carried out in different institutions, however in both instances PKHD1L1-deficient mice were compared to littermate controls. As we have determined that PKHD1L1-defficient bundles are more susceptible to noise-induced damage, the variability in the acoustic environment between the animal facilities experienced by PKHD1L1-defficient mice used in these studies may account for the differences seen in hearing thresholds. Interestingly, audiogram profiles of human patients with non-syndromic hearing loss and biallelic PKHD1L1 variants were also diverse (Redfield et al., 2024). As PKHD1L1 variants may render hair cells more vulnerable to damage due to environmental factors, it is likely that in addition to the potential differences in the damaging effect of distinct PKHD1L1 variants at the protein level, hearing loss profiles in affected probands may also reflect differing levels of acoustic exposure throughout their lives.

With increasing evidence to support PKHD1L1 as a human deafness gene (Lewis et al., 2023; Redfield et al., 2024), the function of PKHD1L1 requires further study. The role of the developmental stereocilia coat remains unclear. Although PKHD1L1 does not localize to any known link structures during its expression period, PKHD1L1 has many protein binding domains. Identification of PKHD1L1’s binding partners would enable the identification of further coat proteins and provide a better understanding of the role of the stereocilia coat during bundle maturation and function. However, PKHD1L1 expression is not limited to the inner ear.

PKHD1L1 has been implicated in or identified as a potential biomarker in several cancers, including but not limited to: thyroid cancer (Zheng et al., 2019), lung adenocarcinoma (Kang et al., 2023; L. Wang et al., 2023), renal cell carcinoma (Yang et al., 2023), skin cutaneous melanoma (Kang et al., 2023), cervical cancer (Zou et al., 2022), squamous cell carcinoma (Song et al., 2021), and breast cancer (Saravia et al., 2019). As described above it has also been identified as a component of the platelet sheddome (Fong et al., 2011). Furthermore, rare pathogenic variants of PKHD1L1 have been linked to conditions such as schizophrenia (Shang et al., 2024), in studies involving human subjects, and anxiety-like traits in rodents (Chitre et al., 2023). We expect that the data, structural modelling, and KO line we present here will benefit and inspire more studies to elucidate the role of PKHD1L1 across other areas of biology.

In conclusion, although the specific molecular function of PKHD1L1 remains elusive, in the present study we build on our current understanding of the role of PKHD1L1 in hearing and deafness. After comprehensive characterization of *Pkhd1l1* gene and protein expression during development and in adult mice, we find PKHD1L1 protein expression on the stereocilia bundle is restricted to early postnatal development. PKHD1L1-deficient mice display no disruption to bundle cohesion or tectorial membrane attachment-crown formation during development. However, progressive bundle disruption and missing stereocilia are later observed in mice deficient in PKHD1L1, with concurrent delayed onset progressive hearing loss. Furthermore, we show that PKHD1L1-deficient mice are susceptible to permanent hearing loss following moderate acoustic overexposure, at noise levels that induce only temporary thresholds shifts in wild-type mice. Here therefore, we propose that PKHD1L1 expression is necessary for the establishment of robust sensory bundles that are required for prolonged hearing function and resilience to noise exposure.

## Materials and Methods

### Generation of Knockout Mice

We transferred *Pkhd1l1^fl/fl^::Atoh1-Cre^+^* mice from Corey Lab at Harvard Medical School to Mass Eye and Ear Animal Care Facility. These mice exhibit PKHD1L1 deficiency selectively in hair cells, triggered by Cre recombinase expression driven by the *Atoh1* promotor. As previously reported (Wu et al., 2019), the *Pkhd1l1^fl/fl^* allele contains 2 loxP sites flanking exon 10, encoding part of the second IPT domain of PKHD1L1. Upon *Atoh1* expression in hair cells, Cre recombinase deletes exon 10 (71 bp), leading to a frameshift and a premature stop codon downstream of the excised region (Wu et al., 2019). The resulting PKHD1L1 peptide is 258 amino acids (including signal peptide), and consists of a single complete non-membrane-bound IPT domain, an incomplete IPT domain, and 11 new amino acids before a the premature stop codon (Fig. 4a).

To generate a constitutive KO model, *Pkhd1l1^fl/fl^* mice were crossed with an EIIa-Cre mouse line (The Jackson Laboratory, B6.FVB-Tg(EIIa-cre)C5379Lmgd/J, #003724) to delete exon 10 from germline. Once homozygous *Pkhd1l1^−/−^* alleles were established, the EIIa-Cre allele was bred out of the colony. Genomic PCR implemented the following primer pairs: Pkhd1l1fl-F (TGA CAC AAC ATA CTG AGC AT) and Pkhd1l1fl-R (GGA AAC TCC TGT TGA AAC AA), yielding a 624 bp fl band and 438 bp WT band; Pkhd1l1KO-F (TGACACAACATACTGAGCAT) and Pkhd1l1KO-R (TTTGAAGACCACACTGAGAG), yielding a 422 bp KO band and 828 bp WT band; EIIA-Cre-F (ATTTGCCTGCATTACCGGTC) and EIIa-Cre-R (ATCAACGTTTTCTTTTCGGA), yielding a 349 bp Cre band.

All procedures were conducted in compliance with ethical regulations approved by the Institutional Animal Care and Use Committee of Mass Eye and Ear.

### RNAscope *in-situ* hybridization labeling and analysis

The RNA labeling procedure was carried out as previously described (Ghosh et al., 2022). Mouse temporal bones were fixed in 4% PFA at room temperature for 3 hours. Cochleae were dissected and the organs of Corti were then placed in cold RNAlater solution in a 96 round bottom well plate and stored at 4°C. Samples were dehydrated by a series of 50, 70, 95, and 100% ethanol washes, each for 5 minutes. Hereafter, each step was followed by washes with milli-Q water. Samples were immersed in 3% H_2_O_2_ (RNAscope® Hydrogen Peroxide, ACD) for 30 minutes at room temperature, incubated in low-pH antigen unmasking solution (Citrate-Based, Vector Laboratories) at 65°C (HybEZII oven, ACD) twice, and incubated in protease plus (RNAscope® Protease Plus, ACD) for 30 minutes at room temperature. Samples were then left in water in 4°C overnight. A custom-ordered *Pkhd1l1* probe (mRNA region 1061-2157bp; exons 12-19) as well as positive and negative control probes were obtained from ACD and applied to samples for 2-hours of hybridization at 40°C. Signals were amplified using the RNAscope 2.5 HD Detection Reagents - RED (warmed to room temperature). AMP1-4 were conducted at 40°C, while AMP 5-6 at room temperature. The duration for AMP1-6 was 30, 15, 30, 15, 25, and 10 minutes respectively, each step followed by 3 low pH antigen unmasking solution washes for 5 minutes each. After amplification, a RED-A and B cocktail at a ratio of 75:1 (shielded from UV light) was applied to the samples for 10 minutes at room temperature, then washed in water extensively. Samples were stained with DAPI (1:2000 dilution in Ca^2+^, Mg^2+^-free HBSS) for 5 minutes and mounted on slides using Prolong Diamond mountant and imaged with a Leica SP8 confocal microscope with a 63x 1.3 NA objective lens.

Images were processed with ImageJ as maximum intensity projections and analyzed using Cellpose (Stringer et al., 2021)to generate a cell mask for each nucleus based on DAPI labeling. Next, resulting ROIs outlining individual cells were transferred to ImageJ. All ROIs corresponding to IHCs and OHCs were used to measure the average intensity values per pixel within each ROI, while the cell masks representing other cells were discarded. Independently, to validate these results, the images were also analyzed manually in ImageJ by outlining 9-12 OHCs or 3-4 IHCs and resulting ROIs were used for average intensity measurements. As there were no substantial differences between Cellpose-generated ROI intensity measurements using the nuclear stain signal, and similar results obtained by manual ROI assignment encompassing entire hair cell bodies, automated and unbiased Cellpose mask generation was favored. Data were plotted and analyzed in GraphPad Prism.

### Immunofluorescence Labeling

Cochleae were dissected in L-15 medium, fixed in 4% formaldehyde (EMS, #15713) in HBSS for 1 hour and washed with Ca^2+^, Mg^2+^-free HBSS. For mice older than P6, fixed cochleae were decalcified in 0.12M EDTA (pH=7.0) for 2 days, washed, then micro-dissected, and permeabilized in 0.5% Triton X-100 (Thermo scientific, #85111) in Ca^2+^, Mg^2+^-free HBSS for 30 minutes. Samples were blocked with 10% goat serum (Jackson ImmunoResearch, #005-000-121) in 0.5% Triton X-100 on Ca^2+^, Mg^2+^-free HBSS for 1 hour and incubated in 1:200, rabbit anti-PKHD1L1 (Novus Bio #NBP2-13765) overnight at 4℃. Samples were washed with Ca^2+^, Mg^2+^-free HBSS and stained for 2 hours in 1:500, Goat anti-Rabbit CF568 (Biotium #20099) and phalloidin CF488 (1:20, Biotium #00042) in the blocking solution. Stereocilin protein labeling was carried out as previously described (Han et al., 2020). Anti-STRC antibody (used 1:50) was kindly provided by Dr. Jinwoong Bok, and previously validated in their study (Han et al., 2020). Following washes, samples were mounted on slides with the Prolong Diamond antifade kit (ThermoFisherScientific # P36961) and imaged on a Leica SP8 confocal microscope using a 63x 1.3NA objective lens.

### Scanning Electron Microscopy

Mouse cochleae were fixed with 2.5% glutaraldehyde in 0.1 M cacodylate buffer (EMS, #15960) supplemented with 2 mM CaCl_2_ for 1.5-2 hours at room temperature, then washed in distilled water (DW). The bony capsule was carefully drilled away with a diamond micro-drill (Ideal Micro Drill, #67-1200A). The organ of Corti was revealed by removing the lateral wall, and the tectorial membrane was removed. For some cochleae the tectorial membranes were also saved and processed for SEM imaging to assess stereocilia imprints.

Samples were rinsed with 0.1 M sodium cacodylate buffer, then stained with 1% Osmium tetroxide(O) (EMS, #19152O) for 1 hour, followed by the treatment with 1% saturated thiocarbohydrazide(T) diluted in DW for 30 minutes. These treatments were repeated, with 3 × 10 min DW washes between steps, using an alternating O-T-O-T-O sequence. Samples were dehydrated in a series of ethanol washes and critical point dried from liquid CO_2_ (Tousimis Autosamdri 815). Next, samples were mounted on aluminum specimen stubs with carbon tape and sputter-coated with 3.0 nm of platinum (Leica EM ACE600) and imaged with a Hitachi S-4700 FE-SEM.

### Immunogold SEM and image analysis

The labeling was carried out as previously described (Ivanchenko et al., 2021). Cochleae from P4, P6, P8 and P10 mice were dissected in L-15 medium, fixed in 4% formaldehyde (PFA, EMS, #15713) in HBSS for 1 hour, then washed with Ca^2+^, Mg^2+^ free HBSS. Samples from mice older than P4 were decalcified in 0.12M EDTA (pH=7.0) for 2 days after fixation prior to micro-dissection. Samples were incubated for 1 hour in blocking solution containing 10% goat serum (Jackson ImmunoResearch, #005-000-121) diluted in Ca^2+^, Mg^2+^-free HBSS, then stained overnight at 4°C in 1:200 rabbit anti-PKHD1L1, (Novus Bio #NBP2-13765) in blocking solution. Samples were washed in Ca^2+^, Mg^2+^ free HBSS and incubated in 12 nm Gold AffiniPure Goat anti-Rabbit secondary antibody (1:30 Jackson ImmunoResearch #111-205-144) overnight at 4°C. After washing steps, samples were post-fixed with 2.5% glutaraldehyde in 0.1 M sodium cacodylate buffer (pH 7.2, EMS, #15960) supplemented with 2 mM CaCl_2_ for 1 hour, then washed with distilled water. Samples were dehydrated, critical point dried and mounted as mentioned above. Dried samples were sputter-coated with Palladium (~4.0 nm) and imaged with Hitachi S-4700 FE-SEM or FEI Helios 660 using insertable backscatter detectors. The number of gold beads per each row of stereocilia was quantified manually. For every age point, 13 stereocilia bundles were analyzed. Data were plotted and analyzed in GraphPad Prism.

### Quantification of missing stereocilia

The number of missing stereocilia was manually quantified from SEM images in three cochlear regions. Data were plotted and analyzed in GraphPad Prism. The number of hair cells (from 1-4 cochlea), used for quantification in each region per genotype is listed in the table below (Table 1) (*Pkhd1l1^fl/fl^::Atoh1-Cre^+^* mice are listed as *Cre^+^*, *Pkhd1l1^fl/fl^* mice are listed as *Cre^−^)*:

**Table 1.** Number of hair cells used for quantification of stereocilia loss.

|  | 6 weeks old |  | 14 weeks old |  | 9 months old |  |
| --- | --- | --- | --- | --- | --- | --- |
|  | *Cre*<sup>+</sup> | *Cre*<sup>-</sup> | *Cre*<sup>+</sup> | *Cre*<sup>-</sup> | *Cre*<sup>+</sup> | *Cre*<sup>-</sup> |
| 16 kHz | 18 | 11 | 6 | 2 | 7 | 5 |
| 22 kHz | 32 | 9 | 12 | 15 | 12 | 12 |
| 32 kHz | 32 | 7 | 17 | 15 | 11 | 11 |

### Protein sequence analysis and prediction of PKHD1L1 cleavage sites

To predict potential cleavage sites in the PKHD1L1 protein sequence, we compared 10 different PKHD1L1 ortholog sequences extracted from the National Center for Biotechnology Information (NCBI) protein database (Sup. Table S1). Species were chosen based on sequence availability and taxonomical diversity. To analyze the complete sequences, each was split into two protein fragments with < 4,000 amino acids and uploaded into the ProP 1.0. online server tool (Duckert et al., 2004). In Sup. Figure 7 the cleavage site prediction for the Human PKHD1L1 is showed comprising the residue p.R3012 to its C-terminal ending residue. The equivalent protein fragments were selected after protein sequence alignment for the other 9 orthologs (See Sup. Table S1 for NCBI residue numbering including signal peptides). Cleavage sites were predicted using the ProP 1.0. default values. This online tool predicts arginine (R) and lysine (K) pro-peptide cleavage sites using an ensemble of neural networks. Scores of > 0.5 indicate that the residue is predicted to be followed by a pro-peptide cleavage site. The higher the score the more confident the prediction.

For sequence identity analysis of the PKHD1L1 cleavage site regions sequences were aligned with the ClustalW algorithm (Larkin et al., 2007) on Geneious using the default parameters and were color-coded by similarity (Kearse et al., 2012).

### PKHD1L1 domain prediction and AlphaFold2 modelling

A consensus domain structure was reached by comparing UniProt and SMART predictions of mouse (*Mm*) PKHD1L1 domains, and SMART prediction of human (*Hs*) PKHD1L1 domains. Protein sequences are reported in supplementary table S1. Signal peptide and transmembrane domain were predicted by UniProt. Domain regions are reported in supplementary table S2. TMEM2-like domains were identified by structural homology, and as reported previously (Hogan et al., 2003; Redfield et al., 2024), at p.2160-2984 and p.3000-3914. MAD-like domain was identified at p.3915-4058, by structural homology with the MAD12 domain of PCDH15 (De-la-Torre et al., 2018).

AlphaFold2 modelling was used to predict the structure of the mouse PKHD1L1 protein region comprising the residues p.3011-4249, harboring the potential cleavage sites and the antibody epitope of NBP2-13765 (p.3670-p.3739). AlphaFold2 simulation was carried out using the ColabFold v1.5.2-patch server using default parameters (Mirdita et al., 2022). Structures were rendered using the VMD software (Humphrey et al., 1996).

### ABR and DPOAE recordings

ABR and DPOAE thresholds were measured using an EPL Acoustic system in an acoustically and electrically shielded soundproof chamber, as described previously (Kujawa & Liberman, 2009). Mice were anesthetized via intraperitoneal injection of 100 mg/kg ketamine and 10 mg/kg xylazine. Artificial tears were applied to protect the animal’s eyes. The outer ear canals were exposed and the tympanic membranes were examined to ensure normal appearance before calibrated earphone placement. Three subdermal needle electrodes were inserted between the ears (reference), behind the pinna (recording), and near the tail (ground). Upon completion, anesthetized animals were placed on a 37°C heating pad for recovery.

Tone-pip ABR stimuli were varied from 4 to 45 kHz, each had a 5 ms duration, 0.5 ms rise-fall time, and of an alternating polarity at 30 Hz. For each pure tone frequency tested, the sound pressure level (SPL) ranged from 20-80 dB. Responses were amplified (10,000x), filtered (0.3-3 kHz), and averaged (512x) by the Cochlear Function Test Suite. ABR thresholds were determined by the lowest SPL at which a recurring 5-peak waveform was observed. Thresholds were initially assessed by the experiment manually, then confirmed by an independent researcher by automated analysis with ABR Peak Analysis software (Eaton-Peabody Laboratories(Suthakar & Liberman, 2019)). Thresholds were averaged from the analysis of 2 independent researchers at any points of disagreement.

DPOAEs were detected in response to 2 simultaneous pure tones, f1 and f2 (f2/f1 = 1.2; f2 was varied from 4 to 45 kHz in half-octave steps and 10-80 dB SPL in 5 dB increments) with the level ratio L1=L2+10. The algorithm to determine the DPOAE threshold for each frequency was an estimate of the crossing point of two curves, 2f1-f2 (dB) and 2f1-f2Nse (dB). No distinction was made between a precisely 80 dB threshold and one that was above the testing limit. All data were plotted and analyzed in GraphPad Prism.

### Acoustic overexposure

Following baseline ABR and DPOAE analysis (carried out as described above), awake 6-week-old mice were placed unrestrained in small cages and exposed to an octave-band noise (8–16 kHz) for 2 hours at 94 dB SPL, which is identified as the highest non-neuropathic noise level associated with temporary threshold shifts (Jensen et al., 2015). Sound exposure was created by a white-noise source, filtered, amplified, and delivered through a JBL compression driver coupled to an exponential horn attached to the top of the exposure booth. Exposure levels were measured in each cage with a 0.25 inch Brüel and Kjær condenser microphone. ABR and DPOAEs were measured 1 day, 2 days, 2 weeks, and 8 weeks after noise exposure. Threshold shifts from baseline were calculated for each mouse at each frequency and time point. Control mice received no noise exposure. All data were plotted and analyzed in GraphPad Prism.

## Acknowledgments

We would like to thank Dr. Jinwoong Bok for providing anti-STRC antibodies; Dr. Bradley Walters, Dr. Sumana Ghosh for RNAscope protocol and guidance, and Dr. M. Charles Liberman and Dr. Brikha Shrestha for critical reading of the manuscript. This work was supported by NIH R01DC020190 (NIDCD) and R01DC017166 (NIDCD) to A.A.I. and the Speech and Hearing Bioscience and Technology Program Training grant T32 DC000038 (NIDCD). The funders had no role in study design, data collection and analysis, decision to publish, or preparation of the manuscript.

## Author contributions

**OSS**: Validation, Formal analysis, Investigation, Writing - Original Draft, Writing - Review & Editing, Visualization.

**RTO**: Validation, Formal analysis, Investigation, Writing - Original Draft, Writing - Review & Editing, Visualization, Project administration.

**CJT**: Validation, Formal analysis, Investigation, Writing - Review & Editing

**XYZ**: Validation, Methodology, Formal analysis, Investigation, Resources, Writing - Review & Editing.

**EH**: Investigation, Writing - Review & Editing.

**PD**: Methodology, Validation, Formal analysis, Investigation, Writing - Review & Editing, Visualization.

**DH:** Methodology, Investigation, Writing - Review & Editing.

**AAI**: Conceptualization, Formal analysis, Investigation, Resources, Writing - Review & Editing, Supervision, Project administration, Funding acquisition.

## Supplemental materials

**Supplemental Figure 1.**
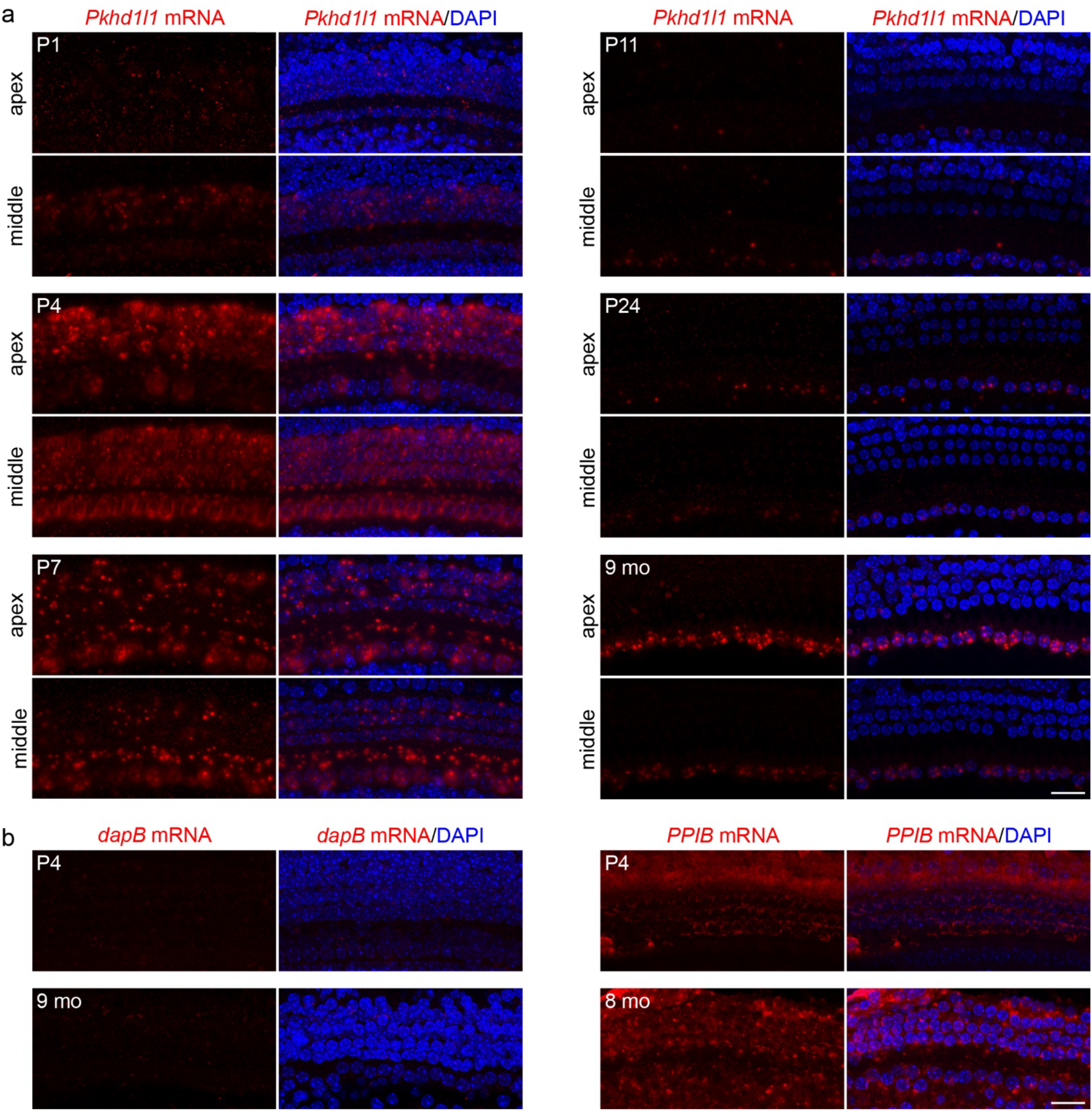
Pkhd1l1 mRNA levels in apical and mid-cochlear hair cells gradually decrease during early postnatal development. **a,** *In situ* detection of *Pkhd1l1* mRNA by RNAscope fluorescence labeling of the apical and mid-cochlear region reveals gradual decrease of mRNA signal in both, OHCs and IHCs by P11, while the mRNA levels in IHCs increase by 9 months. *Red*, *Pkhd1l1* mRNA fluorescence. *Blue*, DAPI. **b,** Negative control: *dapB* mRNA detection (dihydrodipicolinate reductase gene of *Bacillus subtilis*) in the mid-cochlear region shows negligible signal at P4 and 9-month time points. Positive control: *PPIB* mRNA (Cyclophilin B, ubiquitously expressed at low levels providing a rigorous control for sample quality) is detected at comparable levels in the mid-region of P4 and 8-month cochlea. *Scale bars,* 20 µm.

**Supplemental Figure 2.**
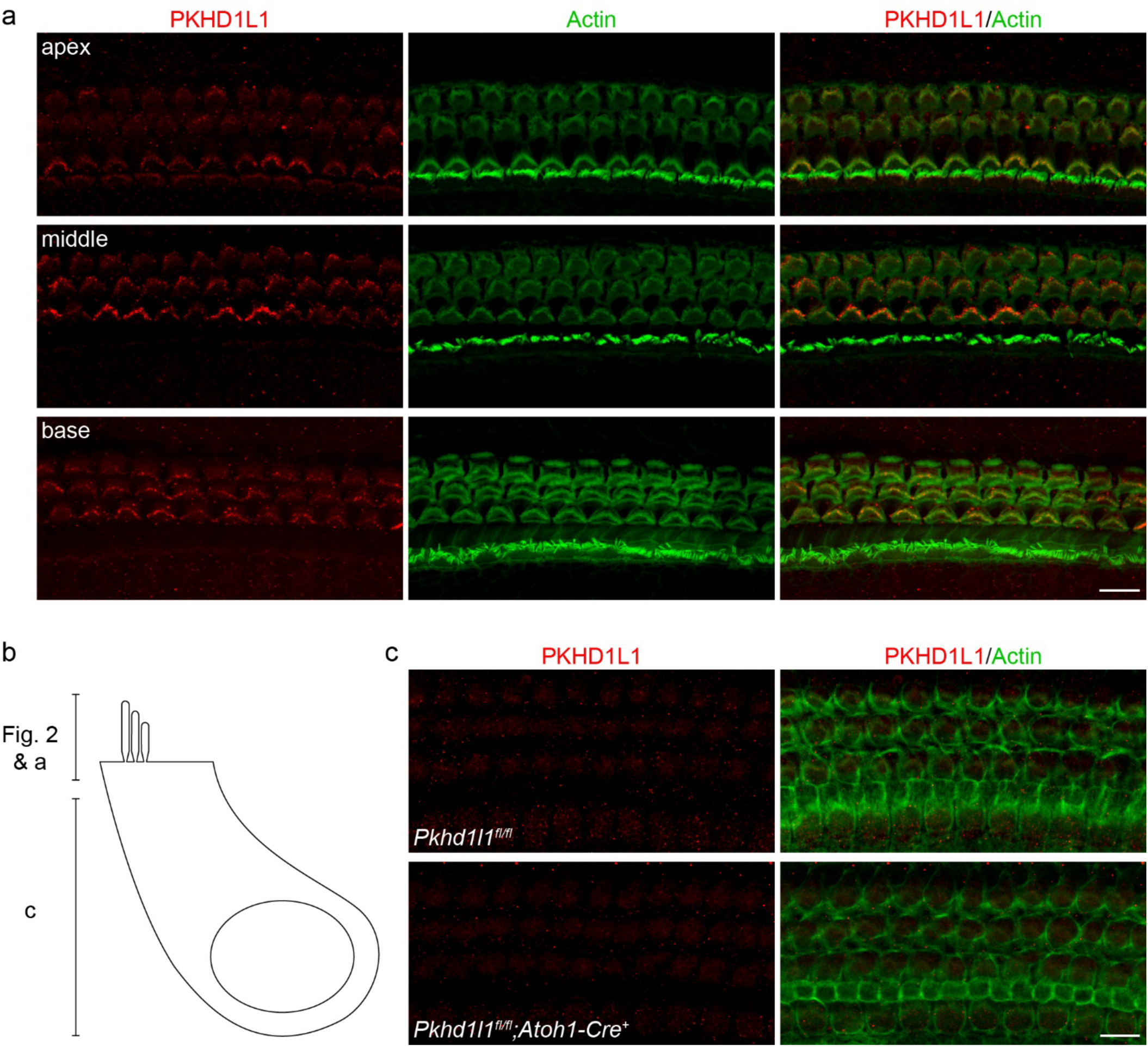
PKHD1L1 is primarily detected within hair-cell stereocilia bundles and not in cell bodies. **a,** Maximum intensity z-projection images of anti-PKHD1L1 labeling present in the hair cell stereocilia at P8 in the apex, middle and basal regions of the cochlea. **b,** Schematic of confocal z-projection depths in Figure 2 and panels a & c. **c,** Maximum intensity z-projection images of anti-PKHD1L1 labeling within the hair cell bodies at P4 in the basal region of the cochlea. No PKHD1L1-specific fluorescence is observed in normal *Pkhd1l1^fl/fl^::Atoh1-Cre^−^* mice compared to *Pkhd1l1^fl/fl^::Atoh1-Cre^+^* negative controls, although some non-specific labeling is present in both genotypes. *Red*, anti-PKHD1L1; *Green*, Phalloidin labeling. *Scale bars,* 10 µm.

**Supplemental Figure 3.**
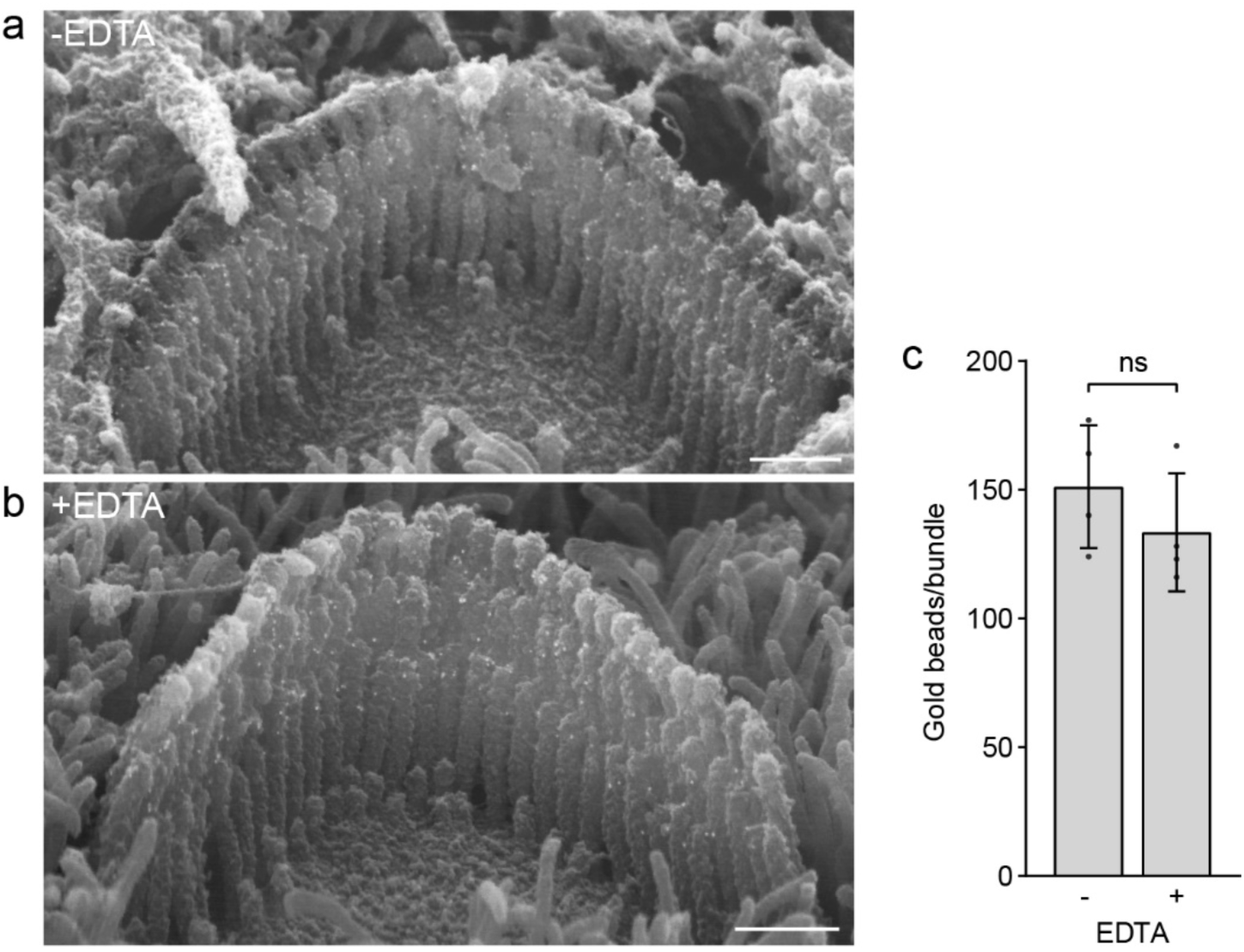
Anti-PKHD1L1 immunogold labeling efficiency is not affected by decalcification. **a**, Representative SEM images of OHC stereocilia bundle labeling of acutely fixed (*top*) and fixed then decalcified (*bottom*) cochlea. *Scale bar*, 500 nm. **b**, Quantification of gold beads per OHC stereocilia bundle. *Statistical analysis*, two-tailed t-test, p>0.05. Data displayed as mean ± SD, points represent individual cells (n=4 in each condition).

**Supplemental Figure 4.**
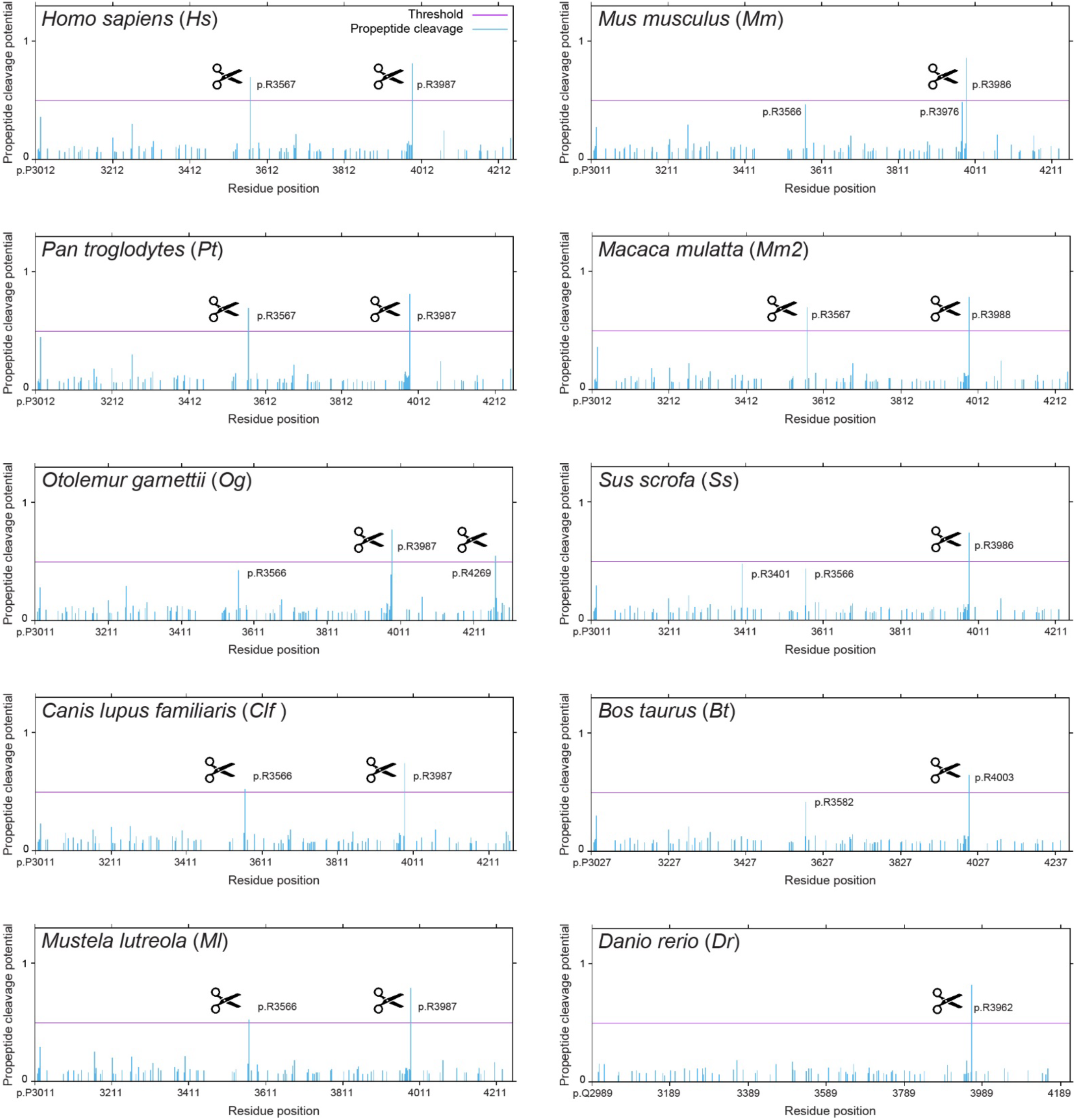
Prediction of potential cleavage sites in PKHD1L1 proteins orthologs. Cleavage sites were predicted across 10 different orthologs for full-length PKHD1L1 protein sequences. Only the region of each PKHD1L1 protein sequence where the cleavage sites are predicted are shown for clarity (see Sup. Table 1 for list of the species and NCBI accession codes). PKHD1L1 protein sequences were analyzed using the ProP v.1.0b ProPeptide Cleavage Site Prediction online server tool. The residue position is indicated for each species, based on NCBI accession numbers (Sup. Table S1) (X axes) and the score for each arginine (R) and lysine (K) residue is shown for the analyzed sequences (Y axes). A score of >0.5 (represented by the purple threshold line) was used to identify residues predicted to be a pro-peptide cleavage site (marked by a black scissor sign). Other plausible cleavage sites below the threshold line are also indicated by residue numbering. A higher score reflects a higher prediction confidence.

**Supplemental Figure 5.**
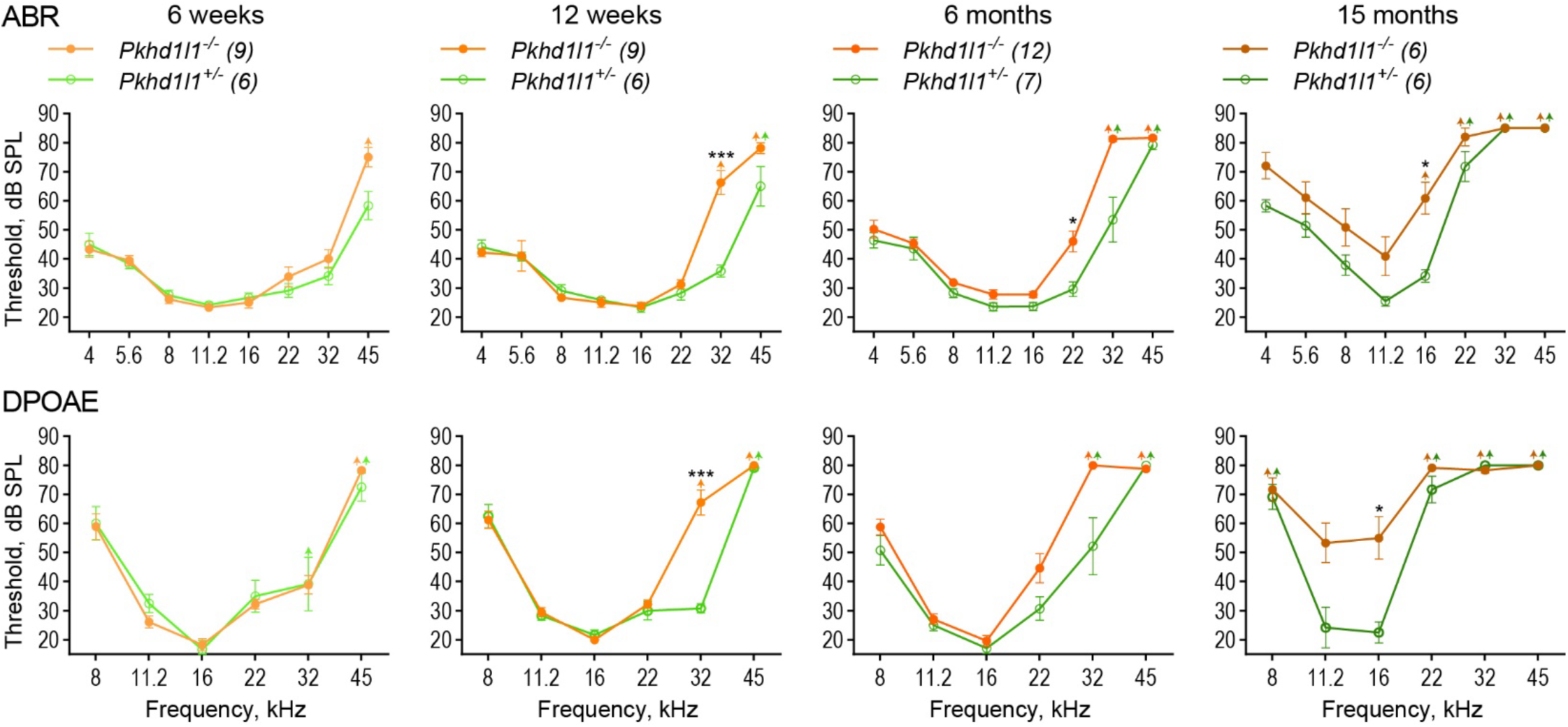
Pkhd1l1^−/−^ KO mice display progressive hearing loss. ABR and DPOAE thresholds of constitutive *Pkhd1l1^−/−^* KO and *Pkhd1l1^+/−^* control littermate mice show progressive high-frequency hearing loss in PKHD1L1-deficient mice as early as 6 weeks. Data are shown as mean ± SEM. Up-arrows indicate that at the highest SPL level tested (80 dB SPL), at least one mouse in the group had no detectable thresholds at that frequency. *Statistical analysis*, two-way ANOVA with matched design between frequencies from the same mouse. Sidak’s multiple comparison tests are shown between genotypes for each frequency. *p<0.05, **p<0.01, ***p<0.001.

**Supplemental Figure 6.**
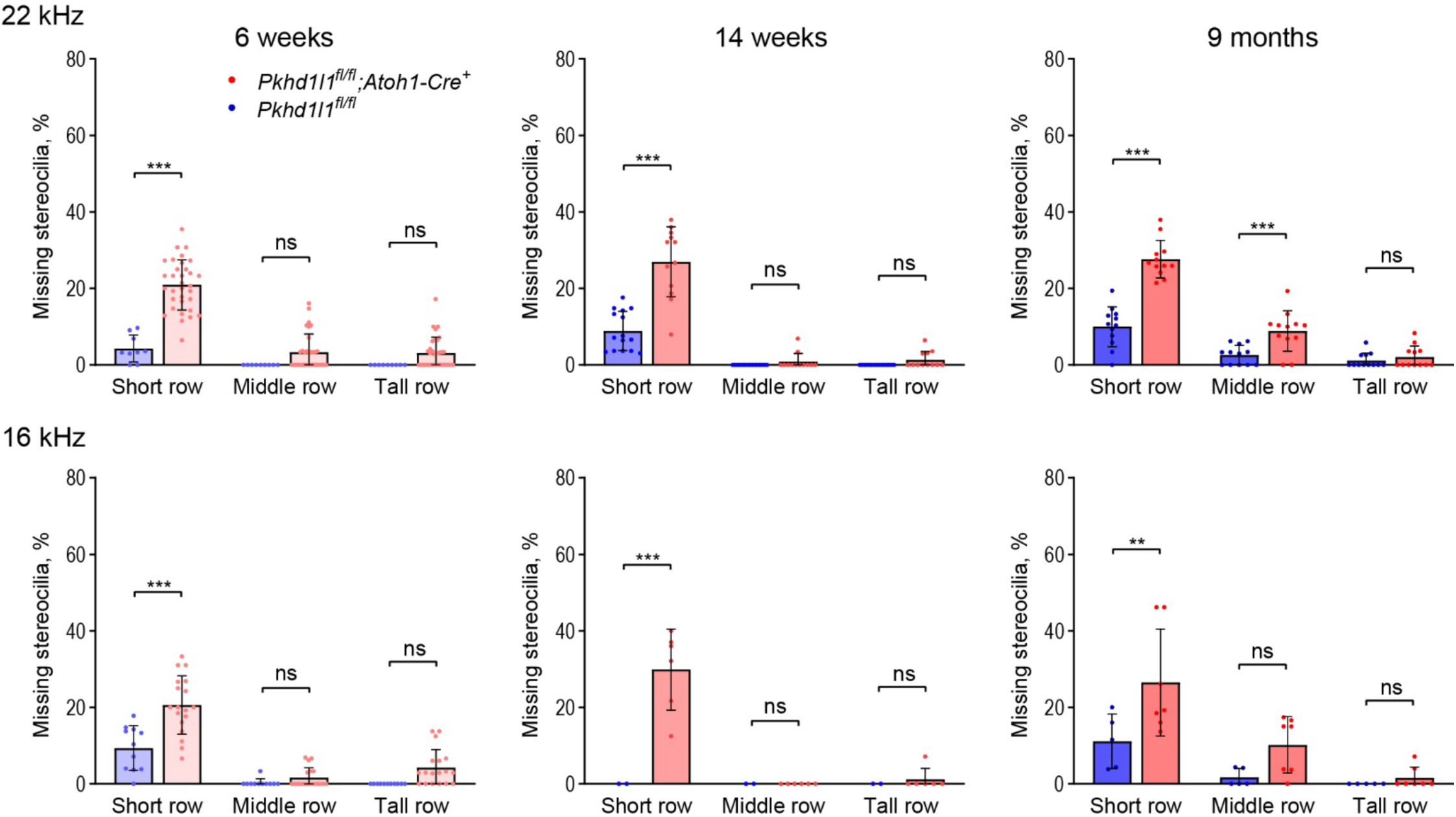
Stereocilia loss in middle and apical regions of the cochlea of PKHD1L1-difficient mice. Quantification of missing stereocilia in OHC bundles in the 22 kHz (middle) and more apical, 16 kHz, regions of the cochlea, presented as percentage of missing stereocilia, per row. There is a greater stereocilia loss in *Pkhd1l1^fl/fl^::Atoh1-Cre^+^* mice as compared to their *Pkhd1l1^fl/fl^::Atoh1-Cre^−^* littermates at all age points. Data displayed as mean ± SD, points represent individual cells (n numbers in methods section). *Statistical analysis*, two-way ANOVA with matched design between rows from the same cell. Sidak’s multiple comparison tests are shown between rows for each genotype. *p<0.05, **p<0.01, ***p<0.001.

**Supplemental Figure 7.**
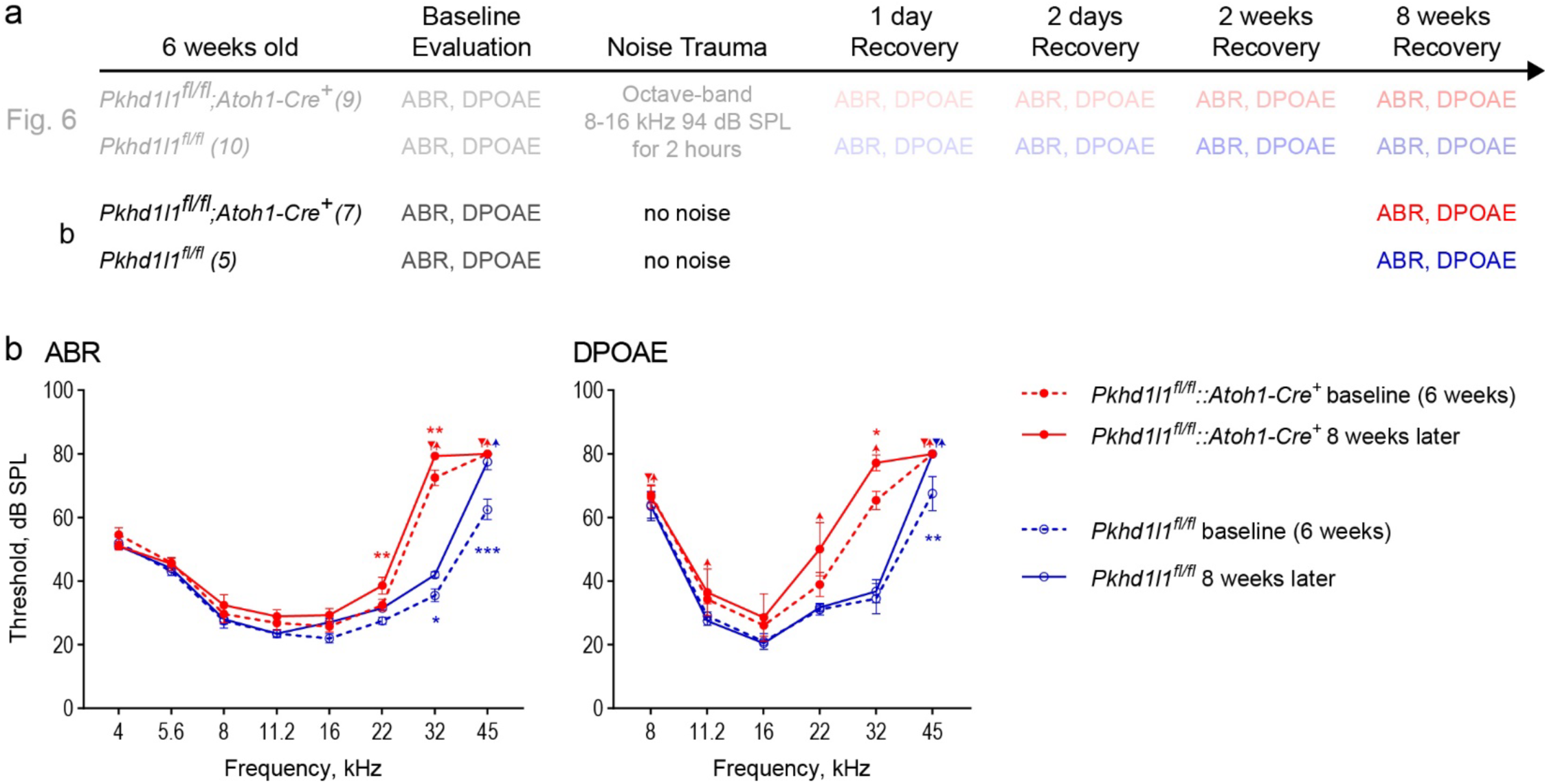
Hearing performance of control mice that underwent no noise trauma exhibit detectable levels of progressive hearing loss 8 weeks later. **a**, Noise exposure experimental design with no noise controls. 6 week old *Pkhd1l1^fl/fl^::Atoh1-Cre^+^* mice and *Pkhd1l1^fl/fl^::Atoh1-Cre^−^* normal mice had baseline ABR and DPOAE evaluation prior to TTS-inducing level of noise trauma (Figure 6), or no noise trauma in controls (b). ABR and DPOAEs were recorded again 8 weeks later. **b**, ABR and DPOAE thresholds in no noise groups at 6 weeks (*dotted lines*) and 8 weeks later (*solid lines*). Both *Pkhd1l1^fl/fl^::Atoh1-Cre^−^* control mice (*blue*) and *Pkhd1l1^fl/fl^::Atoh1-Cre^+^* mice (*red*) show an increase in high frequency thresholds as a result of aging, in addition to the raised thresholds demonstrated in PKHD1L1-deficient mice. Data are shown as mean ± SEM. Inverted triangle (baseline) and up-arrows (8 weeks later) indicate that at the highest SPL level tested (80 dB SPL), at least one mouse in the group had no detectable thresholds at that frequency. *Statistical analysis*, two-way ANOVA with matched design between frequencies from the same mouse, and repeated measures at multiple time points. Sidak’s multiple comparison tests are shown between time points for each frequency for both genotypes. *p<0.05, **p<0.01, ***p<0.001.

**Table S1.**
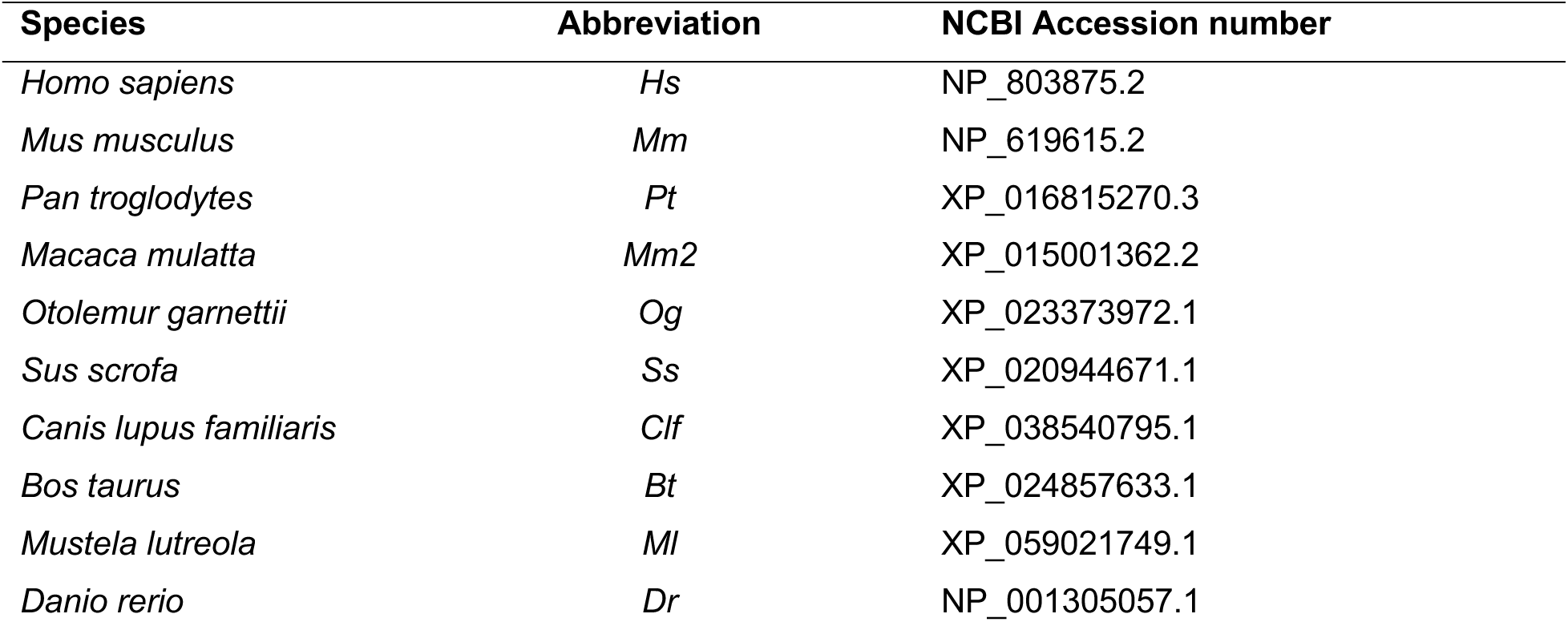
Species and PKHD1L1 accession numbers used for multiple sequence alignment analysis, cleavage site prediction, and AlphaFold2 modelling.

**Table S2.** PKHD1L1 domain predictions.

| Consensus |  |  |  | Mouse UniProt |  |  |  | Mouse SMART |  |  |  | Human SMART |  |  |  |
| --- | --- | --- | --- | --- | --- | --- | --- | --- | --- | --- | --- | --- | --- | --- | --- |
| Domain | Residues |  |  | Domain | Residues |  |  | Domain | Residues |  |  | Domain | Residues |  |  |
| SP | 1 | - | 20 | SP | 1 | - | 20 |  |  |  |  |  |  |  |  |
| IPT | 31 | - | 132 | IPT | 31 | - | 132 | IPT | 30 | - | 141 | IPT | 30 | - | 130 |
| IPT | 146 | - | 255 | IPT | 146 | - | 255 |  |  |  |  | IPT | 145 | - | 257 |
| IPT | 270 | - | 361 | IPT | 270 | - | 361 | IPT | 269 | - | 362 | IPT | 271 | - | 362 |
| PbH1 | 398 | - | 420 | PA14 | 337 | - | 492 | PbH1 | 398 | - | 420 | PbH1 | 398 | - | 420 |
| IPT | 1067 | - | 1153 | IPT | 1067 | - | 1153 | IPT | 1066 | - | 1154 | IPT | 1066 | - | 1152 |
| IPT | 1155 | - | 1234 | IPT | 1155 | - | 1234 | IPT | 1156 | - | 1235 | IPT | 1154 | - | 1235 |
| IPT | 1240 | - | 1323 | IPT | 1240 | - | 1323 |  |  |  |  | IPT | 1239 | - | 1327 |
| IPT | 1329 | - | 1468 | IPT | 1329 | - | 1468 | IPT | 1328 | - | 1407 | IPT | 1329 | - | 1408 |
| IPT | 1565 | - | 1648 | IPT | 1565 | - | 1648 |  |  |  |  | IPT | 1565 | - | 1650 |
| IPT | 1658 | - | 1742 | IPT | 1658 | - | 1742 | IPT | 1657 | - | 1743 | IPT | 1658 | - | 1744 |
| IPT | 1748 | - | 1827 | IPT | 1748 | - | 1827 |  |  |  |  | IPT | 1748 | - | 1829 |
| IPT | 1830 | - | 1909 | IPT | 1830 | - | 1909 | IPT | 1829 | - | 1910 | IPT | 1830 | - | 1911 |
| IPT | 1915 | - | 1996 | IPT | 1915 | - | 1996 | IPT | 1914 | - | 1997 | IPT | 1915 | - | 1998 |
| IPT | 1998 | - | 2084 | IPT | 1998 | - | 2084 | IPT | 1998 | - | 2085 | IPT | 1999 | - | 2086 |
| IPT | 2090 | - | 2175 | IPT | 2090 | - | 2175 | IPT | 2089 | - | 2176 | IPT | 2090 | - | 2177 |
|  |  |  |  |  |  |  |  | PbH1 | 2105 | - | 2126 |  |  |  |  |
| G8 | 2183 | - | 2303 | G8 | 2183 | - | 2303 | G8 | 2183 | - | 2303 | G8 | 2184 | - | 2304 |
| PbH1 | 2484 | - | 2506 | PbH1 | 2484 | - | 2506 | PbH1 | 2484 | - | 2506 |  |  |  |  |
| PbH1 | 2507 | - | 2529 | PbH1 | 2507 | - | 2529 | PbH1 | 2507 | - | 2529 | PbH1 | 2508 | - | 2530 |
| PbH1 | 2565 | - | 2587 | PbH1 | 2565 | - | 2587 | PbH1 | 2565 | - | 2587 | PbH1 | 2566 | - | 2588 |
| PbH1 | 2664 | - | 2686 | PbH1 | 2664 | - | 2686 | PbH1 | 2664 | - | 2686 | PbH1 | 2665 | - | 2687 |
| PbH1 | 2732 | - | 2755 | PbH1 | 2732 | - | 2755 | PbH1 | 2732 | - | 2755 | PbH1 | 2733 | - | 2756 |
| G8 | 3035 | - | 3173 | G8 | 3035 | - | 3173 | G8 | 3035 | - | 3173 | G8 | 3036 | - | 3174 |
| PbH1 | 3292 | - | 3314 | PbH1 | 3292 | - | 3314 | PbH1 | 3292 | - | 3314 | PbH1 | 3293 | - | 3315 |
| PbH1 | 3354 | - | 3376 |  |  |  |  | PbH1 | 3354 | - | 3376 | PbH1 | 3355 | - | 3377 |
| PbH1 | 3415 | - | 3437 | PbH1 | 3415 | - | 3437 | PbH1 | 3415 | - | 3437 | PbH1 | 3416 | - | 3438 |
| PbH1 | 3470 | - | 3492 | PbH1 | 3470 | - | 3492 | PbH1 | 3470 | - | 3492 | PbH1 | 3471 | - | 3493 |
| PbH1 | 3493 | - | 3514 | PbH1 | 3493 | - | 3514 | PbH1 | 3493 | - | 3514 | PbH1 | 3527 | - | 3548 |
| TM | 4223 | - | 4243 | TM | 4223 | - | 4243 |  |  |  |  |  |  |  |  |

